# Notch-Induced Endoplasmic Reticulum-Associated Degradation Governs Thymocyte β−Selection

**DOI:** 10.1101/2021.05.06.443041

**Authors:** Xia Liu, Jingjing Yu, Longyong Xu, Katharine Umphred-Wilson, Fanglue Peng, Yao Ding, Brendan M Barton, Xiangdong Lv, Michael Y Zhao, Shengyi Sun, Yuning Hong, Ling Qi, Stanley Adoro, Xi Chen

## Abstract

Signals from the pre-T cell receptor and Notch coordinately instruct *β*-selection of CD4^−^CD8^−^ double negative (DN) thymocytes to generate *αβ* T cells in the thymus. However, how these signals ensure a high-fidelity proteome and safeguard the clonal diversification of the pre-selection TCR repertoire given the considerable translational activity imposed by *β*-selection is largely unknown. Here, we identify the endoplasmic reticulum (ER)-associated degradation (ERAD) machinery as a critical proteostasis checkpoint during *β*-selection. Expression of the SEL1L-HRD1 complex, the most conserved branch of ERAD, is directly regulated by the transcriptional activity of the Notch intracellular domain. Deletion of *Sel1l* impaired DN3 to DN4 thymocyte transition and severely impaired *αβ* T cell development. Mechanistically, *Sel1l* deficiency induced unresolved ER stress that triggered thymocyte apoptosis through the PERK pathway. Accordingly, genetically inactivating PERK rescued T cell development from *Sel1l*- deficient thymocytes. Our study reveals a critical developmental signal controlled proteostasis mechanism that enforces T cell development to ensure a healthy adaptive immunity.

## Introduction

T cells develop from bone marrow-derived early T-cell progenitors (ETP) through a series of well-orchestrated proliferation and differentiation steps in the thymus. In response to intrathymic interleukin (IL)-7 and Kit ligand, ETPs proliferate and differentiate into CD4^-^CD8^-^ double negative (DN) thymocytes (Freeden-Jeffry et al., 1997). Subsequent differentiation of DN thymocytes into CD4^+^CD8^+^ (double positive, DP) thymocytes depends on whether DN3 stage (CD44^-^CD25^+^) thymocytes successfully undergo “*b*-selection”, the first major checkpoint during αβ T cell development (Shah and Zúñiga-Pflücker, 2014). *b*-selection is initiated by signals from the pre-TCR (a heterodimer of the invariant pre-T*a* and TCR*b* proteins) in DN3 thymocytes that have productively undergone V(D)J recombination at the *Tcrb* locus (Mallick et al., 1993; Michie and Zúñiga-Pflücker, 2002). In addition to cell autonomous signal through the pre-TCR, β-selection also requires signal from the Notch receptor (Ciofani and Zúñiga-Pflücker, 2005; Sambandam et al., 2005). Coordinately, pre-TCR and Notch signals induce DN3 thymocytes to undergo 100-200 fold clonal expansion (Yamasaki et al., 2006; Zhao et al., 2019) as they differentiate into DN4 (CD44^-^CD25^-^) cells which give rise to the DP thymocyte precursors of mature *αβ* T cells. This proliferative burst is crucial for the diversification of the pre-selection TCR repertoire (Kreslavsky et al., 2012) and so must be robustly buffered to ensure adequate number of thymocytes audition for positive selection. *β*-selection imposes a considerable demand for new protein synthesis of the newly rearranged *Tcrb* gene and the multiple factors that execute the transcriptional and metabolic programs demanded by DN thymocyte proliferation. However, how proteome homeostasis or “proteostasis” is regulated during thymocyte development is largely unknown.

Endoplasmic reticulum (ER) is the major subcellular site for synthesis and maturation of all transmembrane and secreted proteins. Protein folding is an inherently error-prone process and is tightly regulated by a myriad of chaperones and enzymes (Balchin et al., 2016; Cox et al., 2018). To maintain proteostasis and normal cell function, cells have evolved highly sensitive and sophisticated quality control systems to ensure the fidelity of protein structure, which is especially important for thymocytes undergoing *β*-selection that must repair protein damage and generate a functional and diverse repertoire of T cell receptors with high fidelity (Feige et al., 2015; Feige and Hendershot, 2013). Two such systems conserved across different species are ER-associated degradation (ERAD) and the unfolded protein response (UPR) (**Figure 1 - figure supplement 1A**) (Brodsky, 2012; Hwang and Qi, 2018; Walter and Ron, 2011). ERAD is the principal protein quality control mechanism responsible for targeting misfolded proteins in the ER for cytosolic proteasomal degradation. The E3 ubiquitin ligase HRD1 and its adaptor protein SEL1L, constitute the most conserved branch of ERAD (Brodsky, 2012; Qi et al., 2017; Ruggiano et al., 2014; Sun et al., 2014). SEL1L recruits misfolded proteins bound by ER protein chaperones to the SEL1L-HRD1 complex, through which the misfolded proteins are retrotranslocated into cytosol, ubiquitinated and degraded by the proteasome in the cytosol with the help of CDC48/ p97 (Brodsky, 2012; Nakatsukasa and Brodsky, 2008). Failure to clear the misfolded proteins in the ER activates the UPR (Brodsky, 2012; Hwang and Qi, 2018; Ruggiano et al., 2014). The UPR is a highly conserved, three-pronged pathway that is activated when the rate of cellular protein production exceeds the capacity of the ER to correctly fold and process its protein load or by various intracellular and extracellular stressors that interfere with the protein folding process. This coordinated response is mediated by three ER-localized transmembrane sensors: IRE1*α*, ATF6α, and PERK (Hetz et al., 2011). Under ER stress, IRE1*α* undergoes oligomerization and *trans*-autophosphorylation to activate its RNase domain to induce unconventional splicing of its substrate XBP1 (Walter and Ron, 2011). ER stress also induces PERK-dependent eIF2*a* phosphorylation and subsequent increased cap-independent translation of ATF4 and induction of CHOP (**Figure 1 - figure supplement 1A**) (Hwang and Qi, 2018; Walter and Ron, 2011).

Here, we show that ERAD, but not the UPR, is the master regulator of physiological ER proteostasis in immature DN thymocytes. The ERAD machinery was critically required for successful *β*-selection of DN3 thymocytes and consequently, ERAD deficiency impeded *αβ* T cell development. Intriguingly, ERAD selectively preserves the cellular fitness of αβ, but not γδ T lymphocytes. We found that Notch signaling directly regulates ERAD gene expression to promote the integrity of ER proteostasis during *β*-selection. Activation of ERAD restricts PERK-dependent cell death in DN3 thymocytes during *β*-selection. Genetic inactivation of *Perk* rescued *β*-selection in *Sel1l*-deficient thymocytes.

## Results

### Stringent protein quality control in β-selected thymocytes

To determine translational dynamics in developing thymocytes, we injected wildtype C57BL/6 (WT) mice with O-propargyl puromycin (OP-Puro), a cell-permeable puromycin analog that is incorporated into newly synthesized proteins as a measure of protein synthesis rates (Jose and Signer, 2019; Tong et al., 2020). Animals were euthanized one hour after OP-Puro injection and thymocyte subsets (gated as shown in **Figure 1 - figure supplement 1B, C**) were assessed by flow cytometry for OP-Puro incorporation (**Figure 1A**). Compared to DP and mature single positive thymocytes, DN2 to DN4 thymocytes incorporated the most OP-Puro (**Figure 1B**), a likely reflection of their high metabolic and proliferative activity (Carpenter and Bosselut, 2010; Kreslavsky et al., 2012; Nagelreiter et al., 2018). To assess the relationship between translational activity and proteome quality, we stained WT thymocytes with tetraphenylethene maleimide (TMI), a cell-permeable reagent that only fluoresces when bound to free thiol groups typically exposed on misfolded or unfolded proteins (Chen et al., 2017; Jose et al., 2020). Intriguingly, despite comparable and high protein synthesis rates across DN2 to DN4 thymocytes, we found that DN4 thymocytes displayed markedly lower levels of misfolded/unfolded proteins (**Figure 1C**). These observations suggested that the DN3-to-DN4 transition, which is initiated by *β*- selection, is accompanied by induction of proteome quality control mechanisms.

**Figure 1.**
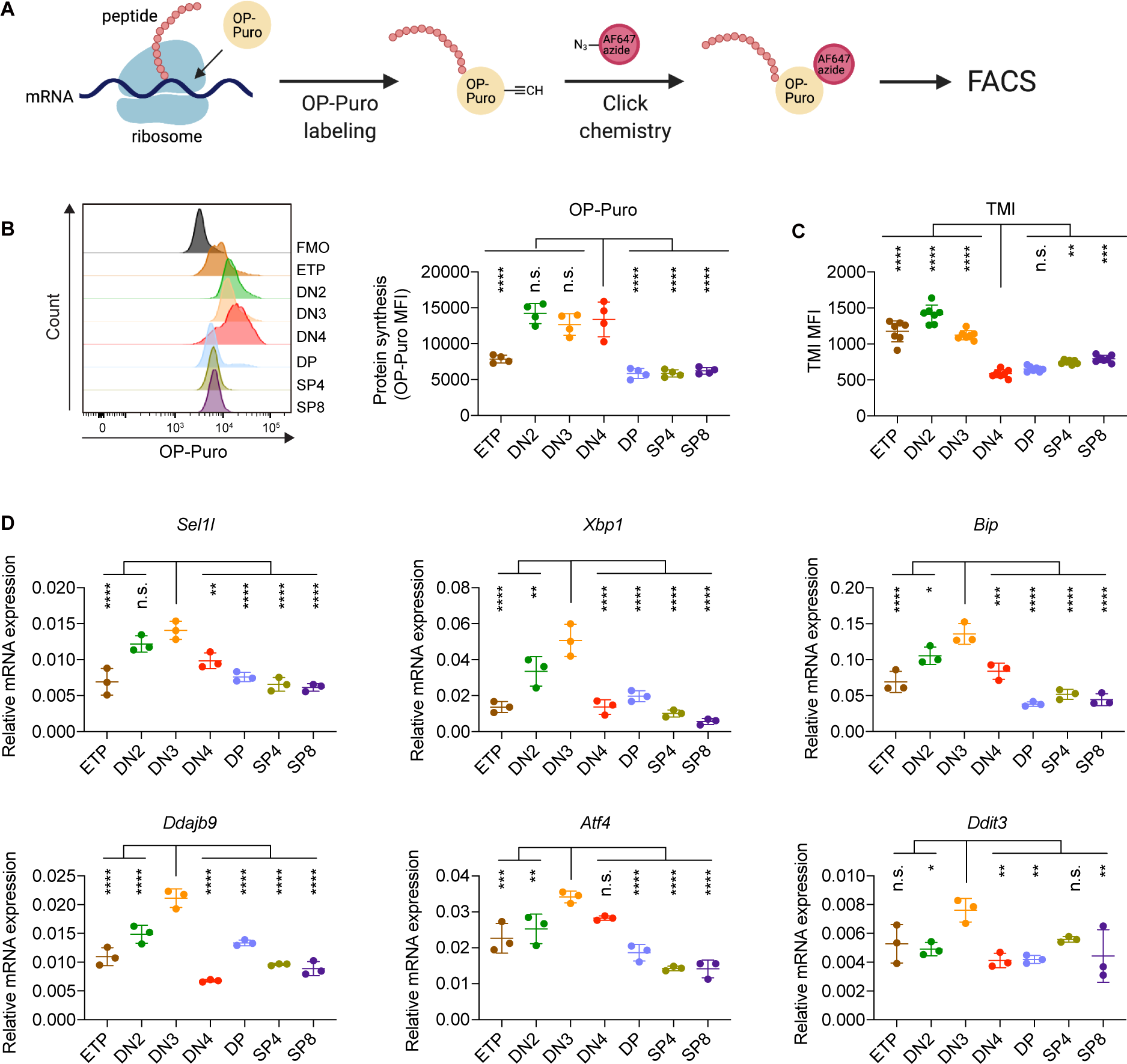
Protein quality control in β-selected thymocytes. **(A)**, Schematic of labeling and detection of nascent protein with OP-Puro. OP-Puro (O-propargyl puromycin) is a cell-permeable puromycin analog that is incorporated into the C-terminus of newly synthesized peptide chain. Fluorophore conjugated with Alexa Fluor 647 was then attached to OP-Puro through a copper-catalyzed click chemistry reaction between alkyne and azide group, which quantifies protein synthesis by fluorescence intensity. **(B),** Representative histogram (**left**) and quantification (**right**) of OP-Puro incorporation in different thymocyte subsets from 8-week-old wild-type mice. FMO represents AF647 control which is the background from the click chemistry in the absence of OP-Puro. MFI, mean fluorescence intensity. *n* = 4 mice. **(C),** Quantification of tetraphenylethene maleimide (TMI) fluorescence in different thymocyte subsets from 8-week-old wild-type mice. *n* = 7 mice. **(D),** Quantitative RT-PCR analysis of ERAD (*Sel1l*) and UPR-related (*Xbp1*, *Bip*, *Dnajb9*, *Ddit3* (*Chop*), *Atf4*) genes expression in different thymocyte subsets from 6-week-old wild-type mice. Data are presented relative to *Actin*; *n* = 3 mice. **(B-D),** ETP: early T lineage precursor (Lin^-^ CD4^-^ CD8^-^ CD44^+^ CD25^-^ CD117^+^); DN2: double negative 2 thymocytes (Lin^-^ CD4^-^ CD8^-^ CD44^+^ CD25^+^); DN3: double negative 3 thymocytes (Lin^-^ CD4^-^ CD8^-^ CD44^-^ CD25^+^); DN4: double negative 4 thymocytes (Lin^-^ CD4^-^ CD8^-^ CD44^-^ CD25^-^); DP: double positive thymocytes (Lin^-^ CD4^+^ CD8^+^); SP4: CD4 single positive thymocytes (Lin^-^ CD4^+^ CD8^-^); SP8: CD8 single positive thymocytes (Lin^-^ CD4^-^ CD8^+^). Results are shown as mean ± s.d. The statistical significance was calculated by one-way ANOVA with Bonferroni test. \**P* < 0.05, \*\**P* < 0.01, \*\*\**P* < 0.001, \*\*\*\**P* < 0.0001, n.s., not significant.

To understand the protein quality control mechanisms operating in thymocytes, we performed quantitative PCR to determine expression of genes encoding ER protein quality control machinery. We found elevated expression of the core ERAD (*Sel1l*) and the UPR (*Xbp1*, *Ddit3 (Chop), Atf4, Dnajb9,* and *Bip*) genes in DN3 thymocytes (**Figure 1D**). Notably, induction of these genes peaked in the DN3 thymocyte stage in which *β*-selection is initiated (Takahama, 2006) and preceded the reduction of misfolded/unfolded proteins in DN4 cells. These results prompted us to hypothesize and explore whether *β*-selection signals induce proteome quality control mechanisms in DN3 cells to enable subsequent stages of thymocyte development.

### The ERAD machinery is required for αβ T cell development

To resolve the ER proteostasis machinery required for the development of thymocytes, we conditionally deleted individual genes encoding key mediators of the UPR (*Xbp1*, *Perk*) and ERAD (*Sel1l*) using the *CD2*-Cre transgene (Siegemund et al., 2015). In this model, significant Cre activity initiated in ETP thymocytes (Shi and Petrie, 2012; Siegemund et al., 2015), and was efficient in depleting genes in subsequent stages of thymocyte development and mature T cells (**Figure 2 - figure supplement 1A, B**). Deletion of the UPR mediator *Xbp1* or *Perk* had no effect on T cell development as thymic cellularity, numbers of DN, DP, single positive (SP) thymocytes and splenic T cell numbers were comparable in gene-deficient and littermate control animals (**Figure 2 - figure supplement 1C-J**). Similarly, *Vav-*Cre*-mediated deletion of Atf6a* which initiates in bone marrow hematopoieitic cell progenitors (Joseph et al., 2013) did not perturb thymocyte development (**Figure 2 - figure supplement 1K-N**).

Strikingly, *CD2*-Cre-mediated deletion of the ERAD core component *Sel1l (Sel1l^CD2^* mice) resulted in a markedly decreased thymus size and cellularity, with significantly reduced and dispersed medullary regions compared to control littermates (*Sel1l^flox/flox^*; Ctrl) (**Figure 2A- D**). The reduced thymus cellularity was accompanied by a profound reduction of peripheral T cells in the spleen and lymph nodes from *Sel1l^CD2^* mice compared to control animal (**Figure 2 - figure supplement 1O-R**). *Sel1l* deletion had no impact on γδ T cells (**Figure 2 - figure supplement 1S**), indicating that within the T-cell lineage, SEL1L is selectively required for αβ T cell development.

**Figure 2.**
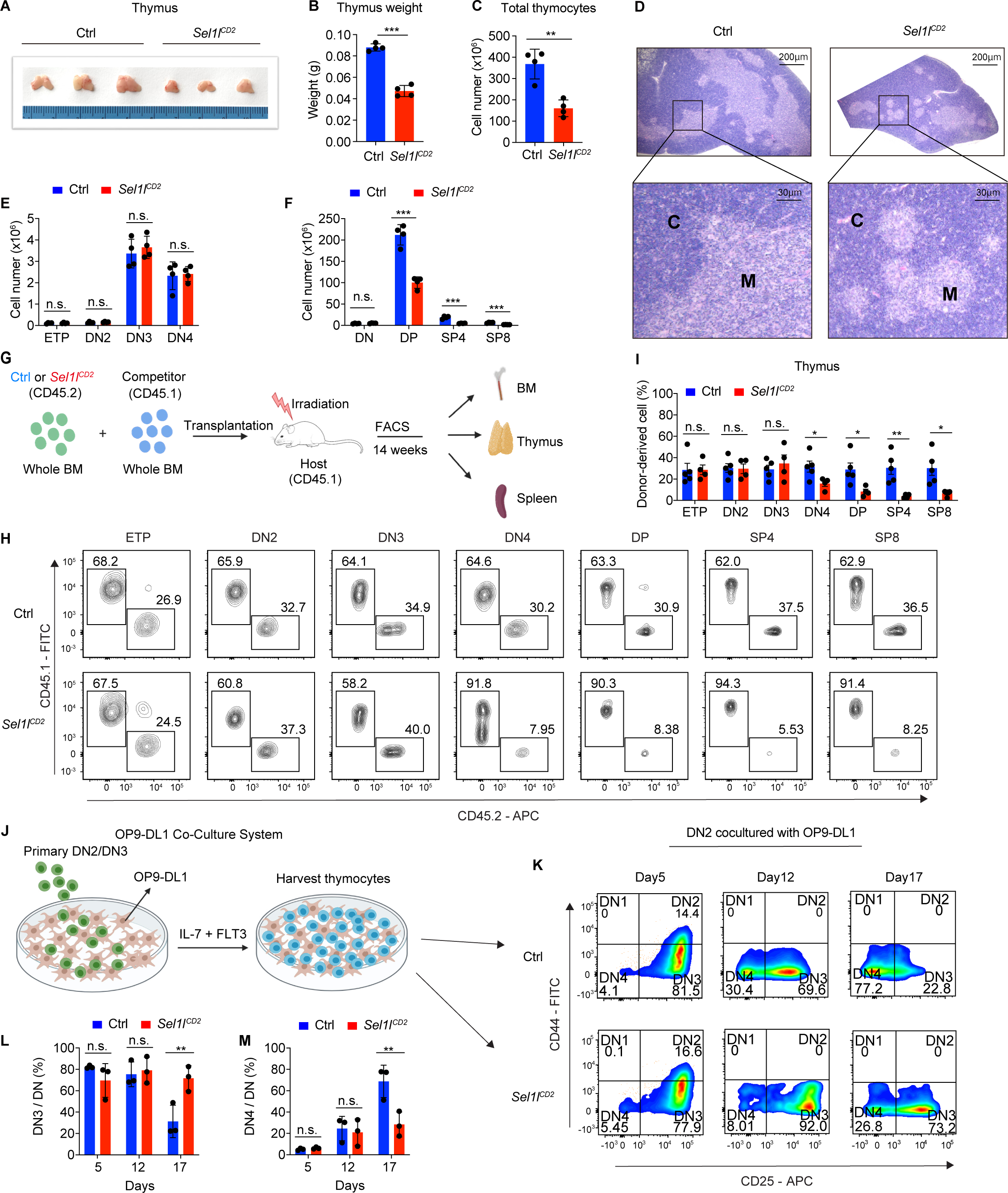
SEL1L is required for αβ T cell development. **(A)**, Images of thymus from 6-8-week-old control (Ctrl, *Sel1l^flox/flox^*) and *Sel1l^CD2^* (*Sel1l^flox/flox^*; *hCD2-iCre*) mice. *n*=3. **(B and C)**, Thymus weight (**B**) and thymus cellularity (**C**) of age and gender-matched control (Ctrl, *Sel1l^flox/flox^*) and *Sel1l^CD2^*-KO (*Sel1l^flox/flox^*; *hCD2-iCre*) mice. *n* =4. **(D)**, Representative images of H&E staining of thymus from 6∼8-week-old control (Ctrl, *Sel1l^flox/flox^*) and *Sel1l^CD2^*-KO (*Sel1l^flox/flox^*; *hCD2-iCre*) mice. Scale bars are indicated. C: Cortex. M: Medulla. **(E and F),** Quantification of cell numbers of the indicated thymocyte subsets in 6-8-week-old control (Ctrl, *Sel1l^flox/flox^*) and *Sel1l^CD2^*-KO (*Sel1l^flox/flox^*; *hCD2-iCre*) mice. *n*=4. **(G)**, Schematic depiction of the competitive bone marrow transplantation (BMT) experiment using whole bone marrow cells from control (Ctrl, *Sel1l^flox/flox^*) or *Sel1l^CD2^*-KO (*Sel1l^flox/flox^*; *hCD2-iCre*) mice as donors. **(H and I),** Representative flow cytometry plots (**H**) and percentage (**I**) of control (Ctrl, *Sel1l^flox/flox^*) or *Sel1l^CD2^*-KO donor-derived thymocyte subsets in the recipient mice 14 weeks after transplantation. *n* = 4-5. **(J),** Schematic overview of OP9-DL1 cell co-culture system. Sorted DN2 or DN3 cells from control (Ctrl) or *Sel1l^CD2^*-KO mice were cultured on a monolayer of OP9 -DL1 cells supplemented with IL-7 and Flt3. (**K**, **L**, **M),** Representative pseudocolor plots (**K**) and percentage of DN3 (**L**) or DN4 (**M**) in DN thymocytes at indicated time points after *in vitro* co-culture of equal number of control (Ctrl, *Sel1l^flox/flox^*) or *Sel1l^CD2^*-KO DN2 cells on OP9-DL1 cells supplemented with IL-7 and Flt3. *n*=3. Results are shown as mean ± s.d. The statistical significance was calculated by two-tailed unpaired t-test (**B, C, E, F, I**) or two-way ANOVA with Bonferroni test (**L, M**). \**P* < 0.05, \*\**P* < 0.01, \*\*\**P* < 0.001, \*\*\*\**P* < 0.0001, n.s., not significant.

### SEL1L is required for DN to DP thymocyte transition following b selection

While *Sel1l^CD2^* mice had similar numbers of DN thymocytes (**Figure 2E** and **Figure 2 - figure supplement 2A**), they showed significantly reduced numbers of DP and mature SP thymocytes compared to littermate controls (**Figure 2F**). This finding suggests that SEL1L functioned during the DN to DP thymocyte transition. To clarify this possibility, we generated and analyzed thymocyte developmental stages in *Sel1l^flox/flox^; CD4-*Cre (*Sel1l^CD4^*) mice. Unlike *CD2*-Cre which initiated in ETP (Siegemund et al., 2015), *CD4*-Cre initiated in immature single-positive (ISP, CD8^+^CD24^+^TCRβ^-^) thymocytes (Gegonne et al., 2018; Kadakia et al., 2019; Xu et al., 2016) and only significantly depleted *Sel1l* in ISPs and later stage thymocytes (**Figure 2 - figure supplement 2B**, **C**). *Sel1l^CD4^* mice exhibited indistinguishable thymic cellularity, immature DN, DP and SP thymocytes numbers from control mice (**Figure 2 - figure supplement 2D-G**). Thus, whereas SEL1L is dispensable for differentiation of post-DN4 thymocytes (i.e., DP, SP and mature T cells), its expression is critical for DN to DP thymocyte differentiation.

To delineate the DN thymocyte developmental stage at which SEL1L is required, we generated 1:1 mixed bone marrow (BM) chimeras by transplanting equal numbers of whole BM cells from control (*Sel1l^flox/flox^*, CD45.2^+^) or *Sel1l^CD2^* (CD45.2^+^) mice along with congenic (CD45.1^+^) wild-type (WT) competitor BM cells into irradiated CD45.1^+^ recipient mice (**Figure 2G**). Fourteen weeks after transplantation, control and *Sel1l^CD2^* donors equally reconstituted similar numbers of all BM hematopoietic progenitors including Lineage^-^Sca-1^+^c-Kit^+^ (LSK) cells, hematopoietic stem progenitor cells (HPC), multipotent progenitor (MPP) and myeloid progenitors (Lineage^-^Sca-1^-^c-Kit^+^ (LS-K) cells (**Figure 2 - figure supplement 2H**). We found substantial defective thymus reconstitution starting from the DN4 stage from *Sel1l*-KO donors (**Figure 2H**, **I**). *Sel1l^CD2^* donor derived DN3 to DN4 ratio was significantly increased compared to controls (**Figure 2 - figure supplement 2I**), suggesting that *Sel1l* depletion compromised the fitness of DN3 thymocytes in which β-selection occurs and impaired their transition to the DN4 stage. The impaired DN thymus reconstitution from *Sel1l^CD2^* donors was accompanied by severe defects in subsequent donor-derived DP and SP thymocytes (**Figure 2H**, **I**), as well as peripheral T cells in the spleen (**Figure 2 - figure supplement 2J**). These data demonstrate a cell-intrinsic requirement of *Sel1l* for DN3 thymocyte progression to later stages of thymocyte differentiation.

To further clarify the requirement for SEL1L in DN3-to-DN4 thymocyte transition during β-selection, we used the *in vitro* T cell differentiation system of culturing immature thymocytes on monolayers of OP9 stromal cells expressing the Notch ligand Delta-like 1 (OP9- DL1 cells) supplemented with IL-7 and Flt3 ligand (Balciunaite et al., 2005; Holmes and Zúñiga-Pflücker, 2009; Schmitt et al., 2004) (**Figure 2J**). We cultured DN2 thymocytes from control or *Sel1l^CD2^* mice on OP9-DL1 cells and found that SEL1L-deficiency markedly abrogated DN4 thymocyte generation (**Figure 2K-M**). On day 17 of co-culture, only 31.3% DN3 cells remained in WT-derived cells, while more than three-fold DN3 cells (71.6%) were found in *Sel1l*-KO derived cells (**Figure 2K-M**), consistent with a block in DN3 to DN4 thymocyte transition. Since only DN3 thymocytes that have successfully undergone β-selection differentiate to the DN4 and DP thymocyte stage these results reinforce that cell-intrinsic SEL1L activity is required for post β-selection DN thymocyte development.

### SEL1L is required for thymocyte survival at the β-selection checkpoint

Next, we assessed the impact of *Sel1l* deletion on DN thymocyte proliferation and survival, two key outcomes of successful *β*-selection (Ciofani and Zúñiga-Pflücker, 2005; Kreslavsky et al., 2012). Whereas BrdU incorporation was similar in control and *Sel1l^CD2^* DN3 thymocytes, we observed more BrdU incorporation in *Sel1l^CD2^* DN4 thymocytes than control DN4 thymocytes **(Figure 3A)**. In agreement, co-staining for Ki-67 and the DNA dye (DAPI) revealed less *Sel1l^CD2^* DN4 thymocytes in G0 phase, and more in G1 and S/G2/M phase **(Figure 3B, C)**.

**Figure 3.**
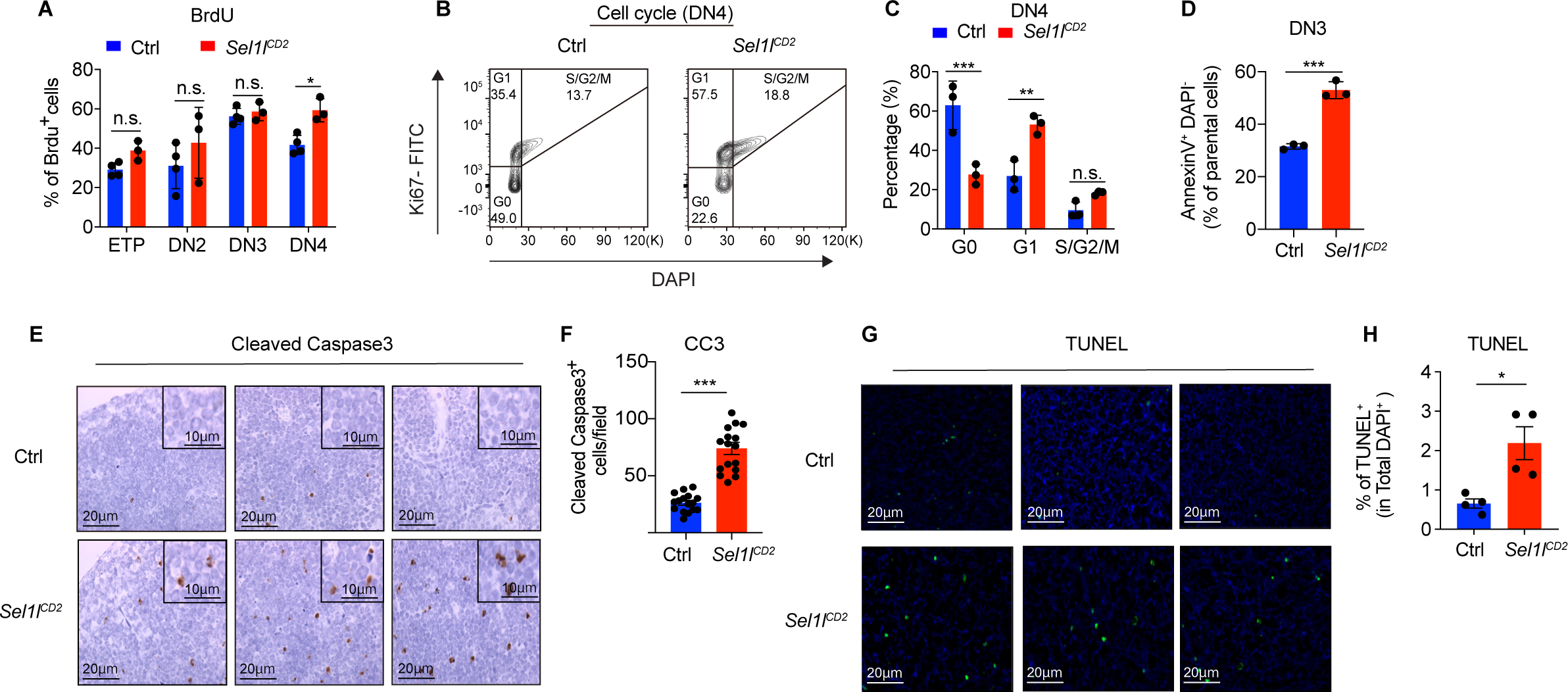
SEL1L is required for thymocyte survival at the β-selection checkpoint. **(A)**, Quantification of BrdU incorporation in different thymocyte subsets from 6-week-old control (Ctrl, *Sel1l^flox/flox^*) or *Sel1l^CD2^-*KO mice. *n*=3-4. **(B and C),** Cell cycle analysis of DN4 thymocytes in 6-week-old control (Ctrl, *Sel1l^flox/flox^*) and *Sel1l^CD2^* mice using Ki67 and DAPI. Representative flow cytometry plots (**B**) and quantification (**C**) are shown. *n*=3. **(D)**, Quantification of apoptotic Ctrl or *Sel1l^CD2^-*KO DN3 thymocytes co-cultured with OP9-DL1 cells *in vitro* for 2 days. *n* = 3. **(E and F)**, Representative images (**E**) and quantification (**F**) of cleaved caspase-3 (CC3) positive cells in the thymus of 6-8-week-old control (Ctrl, *Sel1l^flox/flox^*) or *Sel1l^CD2^*-KO mice. 16 fields were counted at 20× magnification from 4 Ctrl or *Sel1l^CD2^* mice. Scale bars are indicated. **(G and H)**, Representative images (**G**) and quantification (**H**) of TUNEL positive cells in the thymus of 6-8-week-old control (Ctrl, *Sel1l^flox/flox^*) or *Sel1l^CD2^* mice. *n*=4. Scale bar, 20 μM. Results are shown as mean ± s.d. The statistical significance was calculated by two-tailed unpaired t-test (**D, F, H**) or two-way ANOVA with Bonferroni test (**A, C**). \**P* < 0.05, \*\**P* < 0.01, \*\*\**P* < 0.001, \*\*\*\**P* < 0.0001, n.s., not significant.

*Sel1l^CD2^* DN3 cells showed similar cell cycle kinetics as control DN3 thymocytes, in line with their normal levels of BrdU incorporation **(Figure 2 - figure supplement 2K, L)**. The higher proliferation of endogenous *Sel1l*-deficient DN4 thymocytes may be a compensation for their compromised fitness. This likely explains the paradoxical observation that *Sel1l^CD2^* donors generated markedly reduced DN4 thymocytes in BM chimeras (**Figure 2H, I**), yet *Sel1l^CD2^* mice had comparable DN4 thymocytes as control mice in steady state (**Figure 2E**).

That, despite their higher proliferative status, *Sel1l^CD2^* DN4 thymocytes failed to progress to DP thymocytes and generate mature T-cells prompted us to ask if *Sel1l* deficiency resulted in apoptosis of *β*-selected thymocytes. To test this possibility, we cultured equal number of DN3 thymocytes from control or *Sel1l^CD2^* mice on OP9-DL1 stromal cells and assessed their apoptosis *in vitro*. Indeed, *Sel1l^CD2^* DN3 thymocytes showed higher apoptosis measured by proportions of Annexin-V positive cells (**Figure 3D**). To confirm that *Sel1l*-deficiency resulted in DN thymocyte apoptosis *in vivo*, we histologically enumerated apoptosis in thymic sections by measuring active Caspase-3 which degrades multiple cellular proteins and is responsible for morphological changes and DNA fragmentation in cells during apoptosis (Bai et al., 2013). Compared to controls, *Sel1l^CD2^* thymus showed more active Caspase-3 apoptotic cells in the cortex area (**Figure 3E, F**), which is typically populated by DN thymocytes (Trampont et al., 2010). This observation was further confirmed using TUNEL (terminal deoxynucleotidyl transferase dUTP nick end labeling) staining (**Figure 3G, H**). Taken together, we concluded that *Sel1l-*deficient *β*-selected DN3 cells were undergoing apoptosis as they differentiated into DN4 thymocytes.

### Notch directly regulates transcription of ERAD genes

Having established that the SEL1L-ERAD, but not the UPR, is a crucial proteostasis machinery required for *β*-selection, we next sought to understand the thymic signals which regulate SEL1L expression in DN thymocytes. Because high *Sell1l* expression (**Figure 1D**) coincided with high levels of Notch1 in DN2 and DN3 thymocytes (**Figure 4A**), we explored the possibility that Notch ligands might activate the ERAD machinery to enable DN thymocytes to maintain proteostasis during *β*-selection. Stimulation of the thymoma cell line EL4 with Notch ligand Delta-like 4 (DLL4) induced expression of the genes involved in ERAD including *Sel1l*, *Hrd1*, *Os9* and *Edem1* as well as SEL1L proteins (**Figure 4B, C**). Induction of these genes by DLL4 was concomitant with the induction of classical Notch targets like *Hes1*, *Deltex1,* and *Ptcra* (pre- T*α*) (**Figure 4 - figure supplement 1A**).

**Figure 4.**
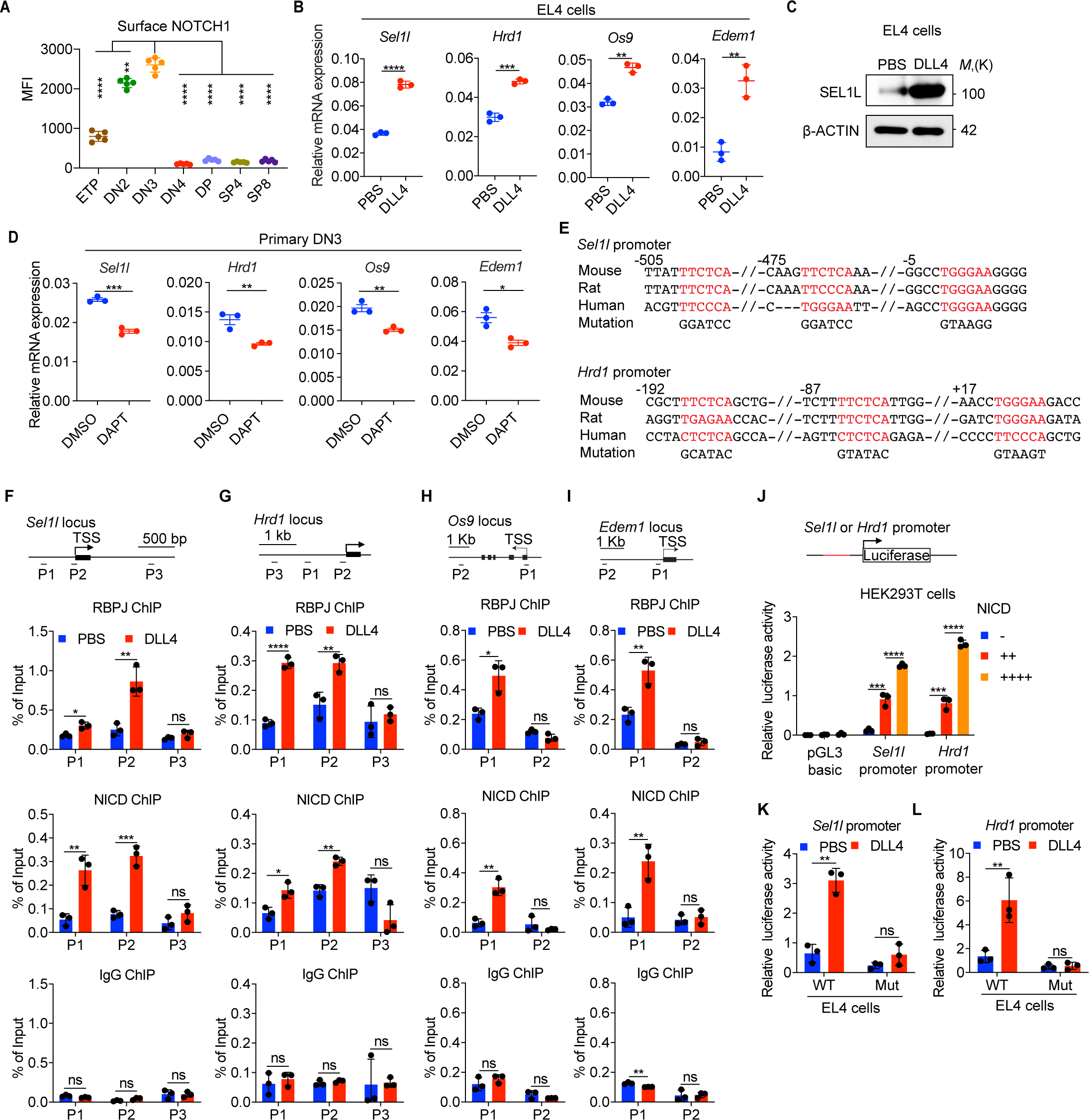
Notch directly regulates transcription of ERAD genes. **(A)**, Quantification of surface NOTCH1 levels in different thymocyte subsets from wild-type mice. MFI, mean fluorescence intensity. *n*=4 mice. **(B)**. Quantitative RT–PCR analysis of ERAD genes (*Sel1l*, *Hrd1*, *Os9*, *Edem1*) expression in EL4 cells after stimulation with 5 μg/ml Delta ligand 4 (DLL4) for 24h. Data are presented relative to *Actin*. *n*=3. **(C)**, Western blot analysis of SEL1L level in EL4 cells after stimulation with Delta ligand 4 (DLL4) for 12h. *β*-ACTIN was used as loading control. The original western blot images are provided in Figure 4**-source data 1**. **(D)**, Quantitative RT–PCR analysis of ERAD genes (*Sel1l*, *Hrd1*, *Os9*, *Edem1*) expression in primary DN3 thymocytes treated with 2mM *γ*-secretase inhibitor DAPT for 5 h. Data are presented relative to *Actin*. *n*=3. **(E)**, Conserved RBP-J binding motif (**Red**) within the promoters of *Sel1l* and *Hrd1*. Alignment of the *Sel1l* (**Upper**) or *Hrd1* (**lower**) promoter from genomic sequence from human, mouse and rat. The numbering corresponds to the mouse sequence and is relative to the transcription start site (TSS). Mutations of the RBP-J binding motifs within *Sel1l* or *Hrd1* promoter luciferase reporters (as in **L**, **M**) are shown. **(F**-**I)**. **Upper**: Schematic diagram of the ChIP primer (P1-P3) locations across the *Sel1l* (**F**), *Hrd1* (**G**), *Edem1* (**H**), or *Os9* (**I**) promoter regions. TSS: transcription start site. **Lower**: Chromatin extracts from EL4 cells treated with PBS or 5 μg/ml DLL4 for 24h were subjected to ChIP using anti-RBP-J antibody, anti-NICD antibody, or normal IgG. Genomic regions of *Sel1l* (**F**), *Hrd1* (**G**), *Edem1* (**H**), or *Os9* (**I**) promoter (as in left panel) were tested for enrichment of RBP-J, NICD or IgG. Data are shown as percentage of input. **(J)**, *Sel1l* or *Hrd1* promoter luciferase reporter was co-transfected with empty vector or different doses of NICD into HEK293T cells, and luciferase activity was measured 36 hours after transfection. pGL3 basic was used as control. **(K and L),** Wild-type or mutant (RBP-J motif mutations, as shown in **e**) *Sel1l* (**k**) or *Hrd1* (**l**) promoter luciferase reporter was transfected into EL4 cells which were treated with PBS or 5 μg/ml DLL4 for 24 hours before harvest. Luciferase activity was measured 36 hours after transfection. All luciferase data are presented relative to *Renilla* readings. Data are shown as mean ± s.d. Two-tailed Student’s t-tests (**A**, **B**, **D, F-I**, **K**, **L**) or one-way ANOVA with Bonferroni test (**J**) were used to calculate *P* values. n.s., not significant, \**P* < 0.05, \*\**P* < 0.01, \*\*\**P* < 0.001, \*\*\*\**P* < 0.0001.

To further determine whether Notch regulates expression of ERAD component genes, we treated freshly isolated primary DN3 thymocytes with the highly specific *γ*-secretase inhibitor DAPT (N-[N-(3,5-difluorophenacetyl)-L-alanyl]-S-phenylglycine t-butyl ester) that blocks ligand-induced cleavage of the Notch intracellular domain (NICD), preventing its nuclear translocation and subsequent transactivation of the RBP-J transcription factor at target genes (Chen et al., 2019; Obaldia et al., 2013; Schmitt et al., 2004), (Bray, 2016). Indeed, treatment with DAPT significantly reduced expression of Notch target genes *Hes1*, *Deltex1,* and *Ptcra* (**Figure 4 - figure supplement 1B**) and also reduced *Sel1l, Hrd1*, *Os9* and *Edem1* expression in DN3 thymocytes (**Figure 4D**). These data demonstrate that Notch signals regulate expression of genes constituting the ERAD machinery.

Analysis of the promoters of the core ERAD genes, *Sel1l* and *Hrd1*, revealed conserved binding sites for RBP-J (**Figure 4E**), the DNA binding partner and master transcription factor of the NICD transactivation complex (Castel et al., 2013; Tanigaki and Honjo, 2007). To interrogate how Notch regulates the ERAD gene components, we performed chromatin immunoprecipitation and qPCR of target DNA (ChIP-qPCR) experiments in EL4 cells. DLL4 stimulation of Notch signaling significantly induced NICD and RBP-J bindings at the promoters of *Sel1l*, *Hrd1*, *Os9* and *Edem* (**Fig. F-I)**.

We cloned the *Sel1l* and *Hrd1* promoters containing the RBP-J binding sites into the pGL3 firefly luciferase reporter and tested the regulation of these promoters by Notch. When co-transfected with promoter luciferase reporters into HEK293T cells, NICD potently induced both *Sel1l* and *Hrd1* promoter activity in a dose-dependent manner (**Figure 4J**). In agreement, DLL4 stimulation of Notch signaling in EL4 cells also substantially activated *Sel1l* and *Hrd1* promoter activity (**Figure 4K, L**). Importantly, mutation of the RBP-J binding sites abolished DLL4- driven induction of *Sel1l* or *Hrd1* luciferase reporter activity (**Figure 4K, L**). These data indicate that Notch signaling directly regulates expression of SEL1L ERAD machinery. Interestingly, although ERAD is known to regulate surface receptor expression in a substrate-specific manner (Boomen and Lehner, 2015; Xu et al., 2020), *Sel1l* deletion did not affect surface Notch1 protein levels in DN thymocyte populations (**Figure 4 - figure supplement 1C**). These results not only suggest that ERAD does not regulate Notch signaling *per se* but also imply that thymocyte *β*- selection defects in *Sel1l^CD2^* mice were not due to failure in Notch signal transduction.

### SEL1L is not required for pre-TCR signaling

The *Tcrb* allele is rearranged in DN2/DN3 thymocytes and the resulting TCRβ protein pairs with pre-Tα and CD3 complex proteins to form the pre-TCR which, together with Notch, transduce *β*- selection signals which promote survival, proliferation and further differentiation of DN3 thymocytes (Ciofani et al., 2004; Michie and Zúñiga-Pflücker, 2002; Sambandam et al., 2005).

DN3 thymocytes that fail to undergo productive recombination of the *Tcrb* locus fail *β*-selection, do not complete the DN4-DP transition and are eliminated by apoptosis (Ciofani et al., 2004).

Therefore, to understand how SEL1L regulates *β*-selection thymocyte survival and the resulting DN-to-DP transition, we first asked whether *Sel1l*-deficiency caused defective V(D)J recombination. Genomic DNA analysis of *Sel1l*-deficient DN3 and DN4 thymocytes showed that recombination of *Vb5-Jb2*, *Vb8-Jb2* and *Vb11-Jb2* gene segments were not altered (**Figure 5 - figure supplement 1A**), indicating that *Sel1l* deficiency does not affect *Tcrb* gene rearrangement.

Next, we asked whether SEL1L regulates the expression and signaling of the pre-TCR complex. Expression of intracellular TCRβ in pre-selected DN3a, post-selected DN3b and DN4 cells was comparable between control and *Sel1l^CD2^* mice (**Figure 5 - figure supplement 1B, C**). *Sel1l* deletion also had no impact on the expression of pre-TCR signaling intermediates including LCK and ZAP-70 in DN3 and DN4 thymocytes (**Figure 5 - figure supplement 1D**).

To further evaluate whether the *Sel1l^CD2^* mice phenotype was due to defective pre-TCR signaling, we introduced the MHCII-restricted TCR-β transgene OT-II into *Sel1l^CD2^* mice. As previously reported (Kim et al., 2014; Marquis et al., 2014), expression of TCR transgenes like OT-II in early DN thymocytes can rescue *β*-selection defects caused by defective pre-TCR expression or signaling. However, the OT-II TCR transgene did not rescue T cell development in OT-II.*Sel1l^CD2^* mice which showed a >80% decrease in thymic cellularity, DP and SP thymocytes (**Figure 5 - figure supplement 1E-G).** In addition, OT-II TCR failed to rescue impaired DN3 to DN4 thymocyte transition resulting from *Sel1l* deficiency as DN4 thymocytes were decreased ∼60% in OT-II.*Sel1l^CD2^* mice compared to littermate controls (**Figure 5 - figure supplement 1H**). Taken together, these data implied that *β*-selection defects in *Sel1l^CD2^* mice were not due to defects in pre-TCR expression or signaling.

### *Sel1l*-deficiency triggers unresolved ER stress during b-selection

To understand the molecular mechanism by which SEL1L ERAD regulates thymocyte survival and differentiation at *β*-selection, we performed RNA-seq on control and *Sel1l^CD2^* DN3 thymocytes. The most upregulated pathways in *Sel1l^CD2^* DN3 thymocytes were ER stress response and the UPR, including both IRE1*α* and PERK pathways (**Figure 5A, B**). Gene set enrichment analysis (GSEA) also revealed enriched ER stress response in *Sel1l^CD2^* thymocytes (**Figure 5C-E**). These signatures hinted at elevated ER stress in *Sel1l^CD2^* DN3 thymocytes.

**Figure 5.**
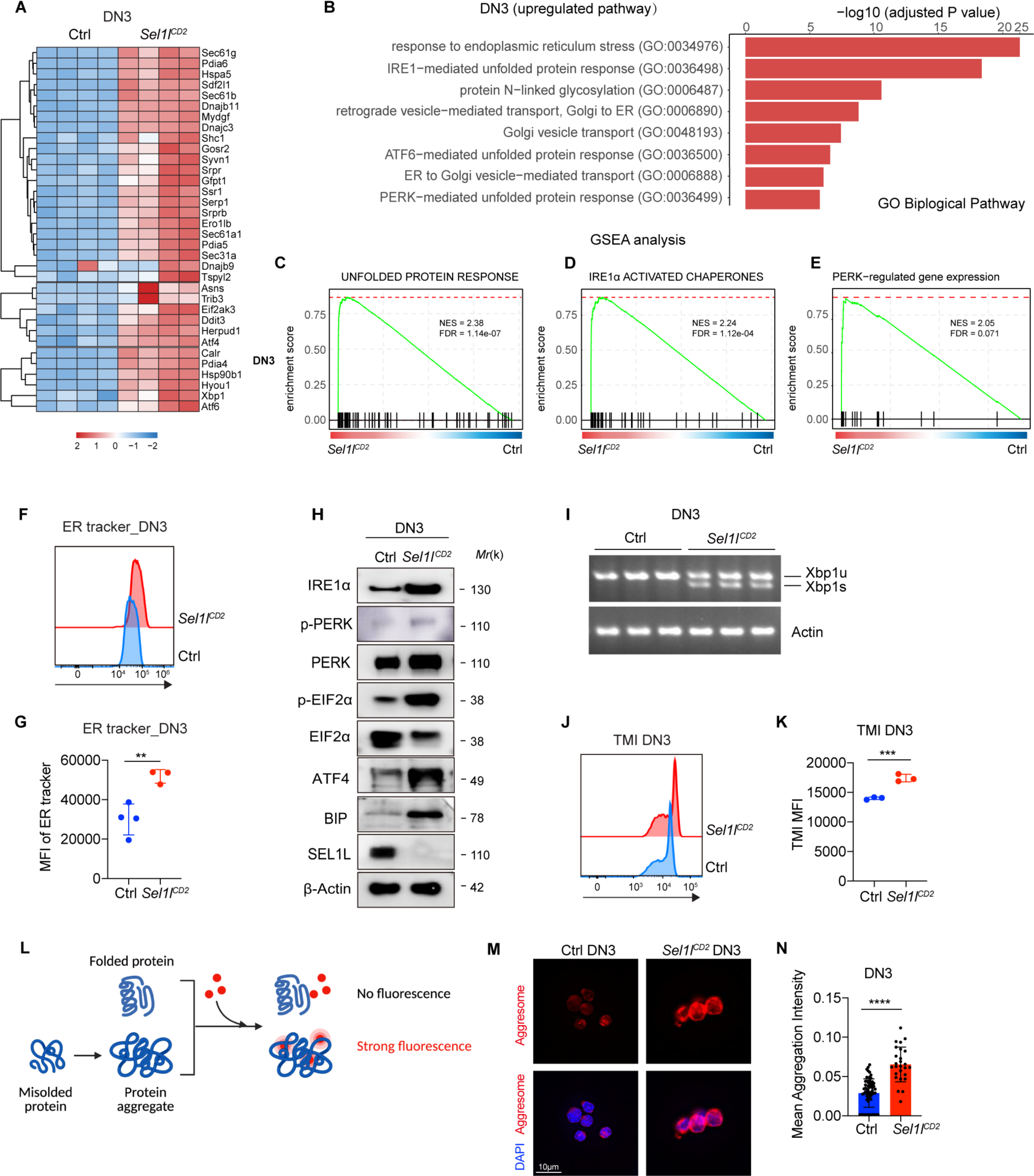
*Sel1l*-deficiency triggers unresolved ER stress during β-selection. **(A)**, Heatmap showing differentially expressed genes from the RNA-seq analysis of DN3 thymocytes sorted from control (Ctrl, *Sel1l^flox/flox^*) or *Sel1l^CD2^*-KO mice. *n*=4. **(B),** Gene Ontology (GO) analysis of the most significantly upregulated pathways in *Sel1l^CD2^*- KO DN3 thymocytes compared with control (Ctrl, *Sel1l^flox/flox^*) DN3 thymocytes. **(C-E),** Plots from GSEA analysis showing enrichment of Unfolded Protein Response (**C**), IRE1*α* (**D**), and PERK (**E**) pathways in *Sel1l^CD2^-*KO DN3 thymocytes compared to control (Ctrl, *Sel1l^flox/flox^*) DN3 thymocytes. **(F and G),** Representative histogram (**F**) and quantification(**G**) of ER-tracker staining in DN3 thymocytes sorted from control (Ctrl, *Sel1l^flox/flox^*) and *Sel1l^CD2^-*KO mice. Ctrl: *n* =4; *Sel1l^CD2^-* KO: *n*=3. MFI, mean fluorescence intensity. **(H)**, Western blot analysis of UPR pathway markers in primary DN3 thymocytes sorted from 6- week-old Ctrl or *Sel1l^CD2^-*KO mice. β-ACTIN was used as loading control. The original western blot images are provided in Figure 5 **- source data 1.** **(I),** PCR analysis of XBP1-splicing in DN3 thymocytes sorted from Ctrl or *Sel1l^CD2^* mice. Xbp1u: Unspliced Xbp1; Xbp1s: Spliced Xbp1. β-ACTIN was used as loading control. The original gel images are provided in Figure 5 **- source data 2.** **(J and K),** Representative histogram (**J**) and quantification (**K**) of unfolded/misfolded protein level measured by TMI in DN3 thymocytes sorted from Ctrl or *Sel1l^CD2^*-KO mice. *n* = 3. **(L)**, Schematic illustration of labeling and detection of misfolded and aggregated proteins with ProteoStat dye. **(M and N),** Representative images (**M**) and quantification (**N**) of protein aggregation measured by ProteoStat Protein Aggregation Detection Kit in primary DN3 thymocytes sorted from 3 pooled Ctrl or *Sel1l^CD2^* mice. Results are shown as mean ± s.d. Two-tailed Student’s t-tests (**G, K, N**) was used to calculate *P* values. \*\**P* < 0.01, \*\*\**P* < 0.001, \*\*\*\**P* < 0.0001.

Consistent with elevated ER stress, flow cytometry quantification of ER tracker dye staining indicated significant ER expansion in *Sel1l^CD2^* DN3 cells compared to control thymocytes (**Figure 5F, G**).

To further ascertain the induction of ER stress following *Sel1l* deletion, we sorted DN3 and DN4 thymocytes from control or *Sel1l^CD2^* mice and examined the activation of all three UPR branches. We found a significant increase in IRE1α proteins and in the splicing of its substrate *Xbp1* in *Sel1l^CD2^* thymocytes (**Figure 5H, I and Figure 5 - figure supplement 2A, B**). We also observed increased PERK and eIF2*α* phosphorylation as well as increased ATF4 and BIP proteins in *Sel1l^CD2^* DN3 thymocytes (**Figure 5H**). Various ER chaperones, including *Calreticulin*, *Grp94 (Hsp90b1)*, *Bip, Hyou1*, and *Canx*, were markedly upregulated in *Sel1l^CD2^* DN3 and DN4 thymocytes (**Figure 5 - figure supplement 2C, D**). These data indicate that *Sel1l* deletion triggers ER stress leading to activation of all three UPR branches in DN thymocytes.

As ERAD alleviates proteotoxic stress by promoting the degradation of misfolded or unfolded proteins we hypothesized that loss of SEL1L increased proteotoxic stress during *β*- selection. Indeed, we found that *Sel1l^CD2^* DN3 thymocytes exhibited significantly higher staining for misfolded/unfolded proteins with TMI (**Figure 5J, K**). We also employed proteostat, a molecular rotor dye, to examine protein aggregation in DN3 thymocytes. The proteostat dye specifically intercalates into the cross-beta spine of quaternary protein structures typically found in misfolded and aggregated proteins, which inhibits the dye’s rotation and leads to a strong fluorescence (**Figure 5L**). *Sel1l^CD2^* DN3 thymocytes displayed more protein aggregates compared with control thymocytes (**Figure 5M, N**). Collectively, these results corroborate that DN thymocytes lacking SEL1L accumulate misfolded/unfolded proteins leading to proteotoxic stress that then trigged the UPR and eventually apoptosis.

### PERK signaling drives b-selected thymocyte apoptosis in *Sel1l^CD2^* mouse

To determine whether upregulation of the UPR contributed to post-*β*-selected thymocyte apoptosis and the *Sel1l^CD2^* mouse phenotype, we generated *Sel1l/Xbp1* double-knockout (*Sel1^flox/flox^.Xbp1^flox/flox^.CD2-cre*), and *Sel1l/Perk* double-knockout (*Sel1^flox/flox^.Perk^flox/flox^.CD2- cre*) mice. Deletion of *Xbp1* in *Sel1l^CD2^* mice did not rescue thymocyte development (**Figure 6 - figure supplement 1A**). In fact, *Sel1l/Xbp1* double-knockout (DKO) mice showed more severe thymocytes development defects including a more than 95% loss in thymus cellularity and DP thymocyte numbers (**Figure 6 - figure supplement 1A**). The DN3 and DN4 thymocytes from *Sel1l/Xbp1* DKO exhibited much more *Chop* expression (**Figure 6 - figure supplement 1B**).

These data suggest that induction of the IRE1*α*/XBP1 pathway functions as a compensatory adaptative pathway to restrain *Sel1l*-deficiency induced ER stress.

In contrast to deletion of *Xbp1*, we found that deletion of *Perk* significantly rescued the *Sel1l^CD2^* mouse phenotype evident in the near complete restoration of thymus cellularity, DP and SP thymocyte cell numbers (**Figure 6A, B**). *Perk* deletion also restored peripheral T cells in spleen and lymph nodes compared to *Sel1l^CD2^* mice (**Figure 6C-F**). Consistent with the rescue, *Perk* deletion significantly reduced *Sel1l*-deficiency induced *Chop* induction (**Figure 6G**) and thymocyte apoptosis (**Figure 6H, I)** and restored normal cell cycle kinetics to *Sel1l^CD2^* DN4 thymocytes (**Figure 6 - figure supplement 1C**). As *Perk* deficiency alone had no effect on T cell development (**Figure 6A-I**), these results indicate that activated PERK signaling contributed to the apoptosis of DN3/DN4 thymocytes that impaired *β*-selection in *Sel1l^CD2^* mice.

**Figure 6.**
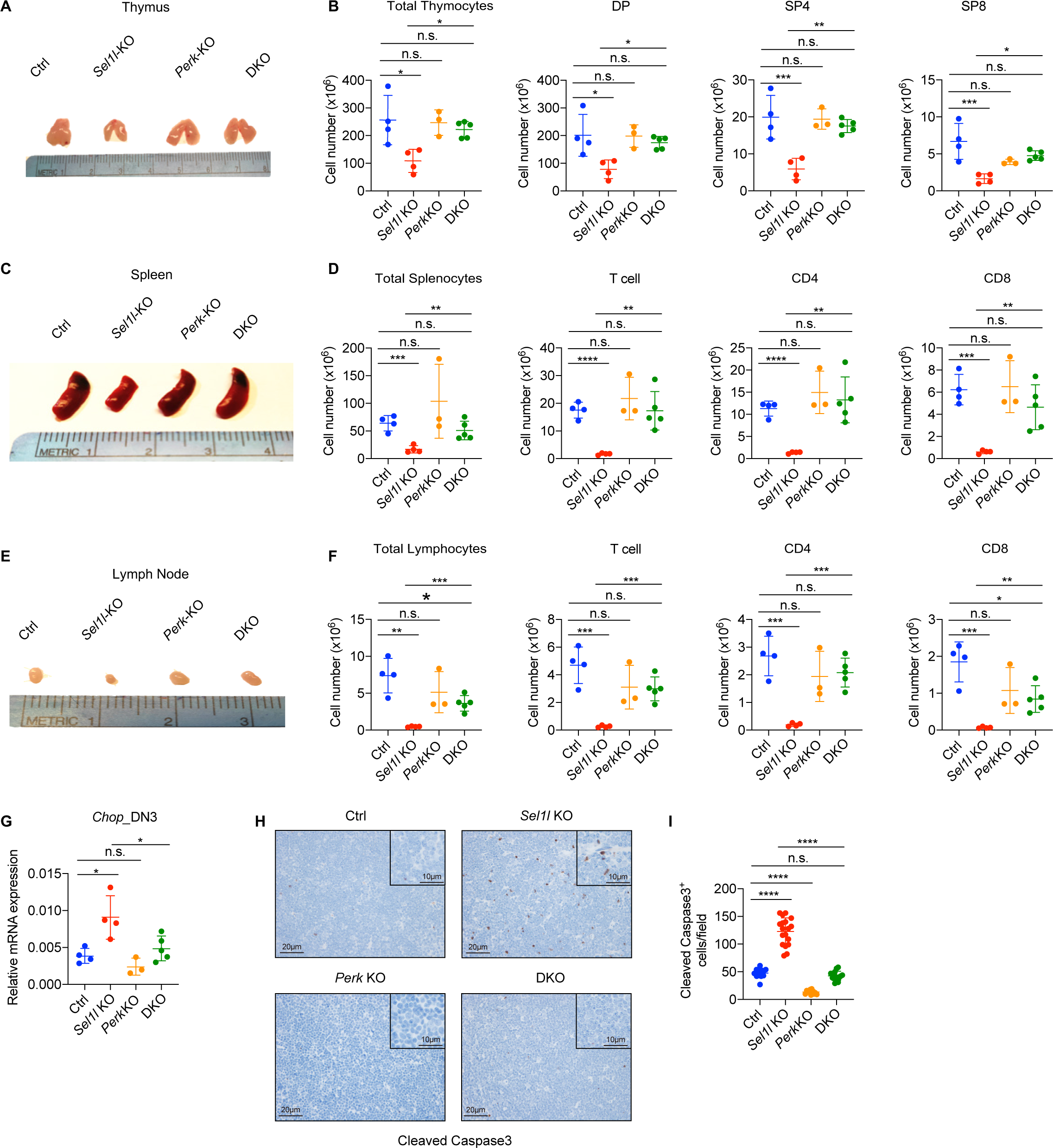
PERK signaling drives β-selected thymocyte apoptosis in *Sel1l^CD2^*-KO mouse. **(A and B),** Representative images of thymus (**A**) and quantification of total thymocytes, DP, SP4 and SP8 thymocytes (**B**) from age (6-week-old) and gender-matched control (Ctrl, *Sel1l^flox/flox^*), *Sel1l*-KO (*Sel1l^flox/flox^* ; *hCD2-iCre*), *Perk-*KO (*Perk^flox/flox^* ; *hCD2-iCre*)and *Sel1l*/*Perk* double knockout (DKO. *Sel1l^flox/flox^, Perk^flox/flox^* ; *hCD2-iCre*) mice. *n* = 3-5 each group. **(C and D),** Representative images of spleen (**C**) and quantification of total splenocytes, total CD3^+^ T cells, CD4^+^ T cells, and CD8^+^ T cells (**D**) from the same mice with indicated genotype as in **A** and **B**. *n* = 3-5 each group. **(E and F),** Representative images of the inguinal (left) lymph node (**E**) and quantification of total lymphocytes, total CD3^+^ T cells, CD4^+^ T cells, and CD8^+^ T cells (**F**) from the same mice with indicated genotype as in **A** and **B**. *n* = 3-5 each group. **(G)**, Quantitative RT–PCR analysis of *Chop* expression in DN3 thymocytes sorted from mice with indicated genotype. *n* = 3-5 each group. **(H and I),** Representative images (**H**) and quantification (**I**) of cleaved caspase-3 (CC3) positive cells in the thymus of 6-8-week-old gender-matched mice with indicated genotype. 12 fields were counted at 20× magnification from 4 mice with indicated genotype. Scale bars are indicated. Data are representative of three independent experiments and are shown as mean ± s.d. The statistical significance was calculated by two-tailed unpaired t-test (**D, F**), One-way ANOVA with turkey test (**B, G**) or one-way ANOVA with Bonferroni test (**I**). ns, not significant, \**P* < 0.05, \*\**P* < 0.01, \*\*\**P* < 0.001, \*\*\*\**P* < 0.0001.

Collectively, we conclude that SEL1L-ERAD promotes *β*-selected DN thymocyte differentiation by maintaining ER proteostasis and suppressing ER stress-induced cell death through the PERK pathway.

## Discussion

In this study, we have uncovered a novel “Notch-ERAD’ axis in thymocyte development. In the absence of the core ERAD protein SEL1L, thymocytes failed to survive *β*-selection and T cell development was severely impaired. Our results imply that the protein synthesis and folding demands during *β*-selection require a robust proteome quality control monitoring which is accomplished by the ERAD in thymocytes transitioning from the DN3 to DN4 stage. Thus, induction of the SEL1L axis of ERAD, which peaks at the DN3 stage, represents a previously undefined and critical ER proteostasis checkpoint during *β*-selection. It is notable that this ER proteostasis checkpoint is also regulated by the same Notch signals which, together with the pre-TCR, induce *β*-selection in DN3 thymocytes. We identified that Notch1 and RBP-J directly bind to most ERAD gene promoters and directly regulate their expression.

ERAD and UPR are two key ER protein quality control machineries that are activated at different thresholds. ERAD is responsible for the clearance of misfolded proteins at steady-state and is constituively activated regardless of ER stress. In contrast, the UPR is a stress response pathway that is triggered when the accumulation of misfolded and unfolded proteins exceed the ER folding capacity. Exisiting evidence suggest that UPR pathways like the IRE1*α*-XBP1 axis appear to selectively regulate hematopoieitic cell differentiation programs but individual UPR enzymes influence early thymopoiesis remain to be fully characterized. For instance, the IRE1*a*- XBP1 is essential for eosinophil (Bettigole et al., 2015), dendritic cell (Cubillos-Ruiz et al., 2015; Iwakoshi et al., 2007; Osorio et al., 2014), NK (Dong et al., 2019; Wang et al., 2019) and plama cell (Reimold et al., 2001; Shapiro-Shelef and Calame, 2005) differentiation. Unlike the marked defect in thymocyte development in the absence of SEL1L-ERAD, our current study now clearly demonstrates that individual UPR regulators are dispensable for *b*-selection and subsequent stages of thymocyte maturation. We specifically found that deletion of the individual UPR master regulators XBP1, ATF6, or PERK had no overt impact on αβ T or γδ T cell development in the thymus and peripheral tissues. Nevertheless, our results don’t exclude the possibility that redundancy might exist among the UPR pathways during thymocyte development.

*Sel1l-*deficiency impaired the survival of DN thymocytes following β selection and resulted in the inability of DN3 thymocytes to expand and progress to DP thymocytes.

Interestingly, this phenotype was not due to defective rearrangement of TCRβ nor defective Notch or pre-TCR signaling which both activate pro-survival genes in *β*-selected thymocytes (Ciofani and Zúñiga-Pflücker, 2005; Kreslavsky et al., 2012; Zhao et al., 2019). Instead, our genetic inactivation delineate a pro-apoptotic pathway driven by the PERK axis. While all three UPR pathways can promote apoptosis and appear to be upregulated in *Sel1l*^CD2^ thymocytes, it is notable that only PERK axis induces apoptosis in *Sel1l*^CD2^ thymocytes. On the contrary, induction of the IRE1*α*-XBP1 axis of the UPR can help cells adapt to stress. Consistent with this view, deletion of *Xbp1* in *Sel1l*^CD2^ thymocytes exacerbated defects in thymocyte development unlike PERK inactivation which restored T cell development from *Sel1l*^CD2^ thymocytes.

Although αβ T and γδ T cells develop in the thymus from the same thymic seeding progenitors, the function of ERAD appears to be restricted to αβ T cells. While future studies are needed to determine the alternative protein quality control mechanism regulating γδ T cell development, our findings are consistent with the selective requirement for Notch in driving *β*- selection (Maillard et al., 2006), a key developmental checkpoint unique to the αβ T cell differentiation program. In addition, DN thymocytes undergoing β selection expand more than 100-fold, a situation that likely explains their increased translational activity. That post-*β*- selected DN thymocytes displayed markedly low levels of misfolded/unfolded proteins support the view that upregulation of the SEL1L-ERAD axis in DN3 thymocytes provides a mechanism to alleviate deleterious proteotoxic stress that compromise the fitness of DN4 thymocytes. In this way, SEL1L-ERAD safegaurds to DN3/DN4 thymocyte pool to ensure that the maximum number of DN thymocytes expressing a functional TCR*β* protein progress to the DP stage to audition for positive selection.

In summary, our study reports a previously unknown function of Notch in maintaining proteostasis and protecting post-*β*-selection thymocytes from ER stress by upregulating ERAD. We propose that in addition to driving energy metabolism and survival programs demanded by proliferating thymocytes following *β*-selection, Notch signals in parallel activate the ERAD machinery in DN3 thymocytes to clear misfolded proteins. Thus, stringent protein quality control through the SEL1L-ERAD pathway is required for successful *β*-selection and the development of the *αβ* T cells that mediate adaptive immunity.

## Acknowledgements

We thank Dr. Juan Carlos Zúñiga-Pflücker (University of Toronto) for providing the OP9-DL1 cells and Dr. Laurie Glimcher (Dana Farber Cancer Institute) for providing the *Xbp1* flox mice. This work was supported by the National Institutes of Health (R01HL146642, R37CA228304 and P50CA186784 to X.C.; R01 AI1143992 and K22CA 218467 to S.A.; R35GM130292 to L.Q.), the US Department of Defense Congressionally Directed Medical Research Programs (W81XWH1910524 to X. C.; W81XWH1910306 to S.A.; W81XWH1910035 to X. L.), and Cancer Prevention and Research Institute of Texas (RP160283 Baylor College of Medicine Comprehensive Cancer Training Program award to F.P.). This work was supported by the Cytometry and Cell Sorting Core at the Baylor College of Medicine with funding from the CPRIT Core Facility Support Award (CPRIT-RP180672), the NIH (CA125123, S10OD025251 and RR024574) and the assistance of J. M. Sederstrom. Imaging for this work was supported by the Integrated Microscopy Core at Baylor College of Medicine and the Center for Advanced Microscopy and Image Informatics (CAMII) with funding from NIH (DK56338, CA125123, ES030285), and CPRIT (RP150578, RP170719), the Dan L. Duncan Comprehensive Cancer Center, and the John S. Dunn Gulf Coast Consortium for Chemical Genomics.

## Author Contributions

X.C. and S.A. conceived the project. X. L, J. Y., L.X., K.U.W., S.A., and X.C. designed the research; X. L., J.Y., L.X., K.U.W., F. P., Y.D., B.M.B., X. L. and M.Y.Z. did the experiments and analyzed the data; S. S., Y.H., and L.Q. contributed to discussions, experimental design and critical reagents; X.C. and S.A. supervised the project; S.A., X.C. and J.Y. wrote the paper.

## Competing interests

The authors have declared that no conflict of interest exists.

## Materials & Correspondence

Correspondence and material requests should be addressed to Stanley Adoro, Ph.D., Department of Pathology, Case Western Reserve University, Cleveland, OH, USA; Phone 216-368-4712; FAX 216-368-0494, or Xi Chen, Ph.D., Department of Molecular and Cellular Biology, Baylor College of Medicine, One Baylor Plaza, MS: BCM130, Debakey Building, BCMM-M626, Houston, TX 77030, USA; Phone 713- 798-4398; FAX 713-790-1275;

## Materials and Method

### Mice

The mice were maintained in a pure C57BL/6 background and kept under specific-pathogen-free conditions in the transgenic mouse facility of the Baylor College of Medicine (22–24 °C, 30– 70% humidity with a 12-h dark and 12-h light cycle). *Sel1l*^flox/flox^ and *Xbp1^flox/flox^* were described previously (Xu et al., 2020)*, Perk^flox/flox^* mice were purchased from Jackson Laboratory (Stock No. 023066). The floxed mice were crossed with either *hCD2-iCre* (The Jackson Laboratory, 008520) or *CD4-Cre* (The Jackson Laboratory, 022071) mice to generate *Sel1l^flox/flox^; hCD2- iCre, Sel1l^flox/flox^; CD4-Cre, Xbp1*^flox/flox^*; hCD2-iCre*, or *Perk*^flox/flox^*; hCD2-iCre* mice.

The *Sel1l^flox/flox^; Xbp1^flox/flox^; hCD2-iCre* mice and *Sel1l^flox/flox^*; *Perk^flox/flox^; hCD2-iCre* mice were generated by crossing *Sel1l^flox/flox^; hCD2-iCre* mice with *Xbp1^flox/flox^* or *Perk^flox/flox^* mice respectively. *Atf6^Vav^-iCre KO* mice were generated by crossing *Atf6l^flox/flox^* (The Jackson Laboratory, 028253) with *Vav-iCre* mice (The Jackson Laboratory, 008610). *Sel1l^CD2^*; OTII transgenic mice were generated by crossing *Sel1l^CD2^* mice with OTII transgenic mice (The Jackson Laboratory, 004194).

Six to Eight-week-old gender-matched mice were used for phenotype analysis and *in vitro* assay. For the bone marrow transplantation assays, female C57BL/6-Ly5.1 (CD45.1^+^) mice (Charles River, 564) were used at 8–12 weeks of age. All procedures were approved by the Baylor College of Medicine Institutional Animal Care and Use Committee or Case Western Reserve University Institutional Animal Care and Use Committee. The study is compliant with all of the relevant ethical regulations regarding animal research.

### In Vivo Assays

For the competitive bone marrow transplantation experiments, CD45.1^+^ recipient mice were lethally irradiated (10 Gy, delivered with two equal doses 4 h apart) and injected retro-orbitally with 1x10^6^ whole bone marrow cells from *Sel1l^flox/flox^* or *Sel1l^flox/flox^*; *hCD2-iCre* mice with an equal number of CD45.1^+^ competitor BM cells. For the BrdU incorporation assay, the mice were injected intraperitoneally with single dose of BrdU (BD, 559619; 50 mg/kg body weight) 2 h before euthanization.

### Flow Cytometry and Cell Sorting

Single-cell suspensions from thymus, spleen and lymph nodes were obtained by passing the tissues through a 70-μm strainer. Bone marrow was removed from femurs and tibiae by flushing with PBS, 2% FBS. For bone marrow and spleen cell suspensions, red blood cells were removed by incubating with RBC Lysis Buffer (Biolegend, 420301). 1 × 10^6^ to 2 × 10^6^ cells were stained with antibodies at 4°C for 30 min in phosphate-buffered saline (PBS) containing 1% bovine serum albumin (BSA), followed by washing to remove the unbound antibodies.

Antibodies used for flow cytometry: The biotin-conjugated lineage markers for excluding non-DN cells are purchased from BioLegend: CD11b (M1/70, 101204), CD11c (N418, 117303), Ter119 (Ter119, 116204), Gr-1 (RB6-8C5, 108404), CD49b (DX5, 108904), and B220 (RA3-6B2, 103204). CD4 (BV650, RM4-5, 100546, BioLegend), CD8 (AF700, 53-6.7, 100730, BioLegend), CD25 (PE-Cy7, 3C7, 101915, BioLegend), CD44 (PE594, IM7, 103056, BioLegend), c-Kit (APC-Cy7, 2B8, 105838, Biolegend), γδTCR (PE, GL3, 118108, Biolegend) and CD27 (APC, LG.3A10, 124211, BioLegend) antibodies were used for analysis of ETP to SP cells in the thymus. CD4 (BV650, RM4-5, 100546, BioLegend), CD8 (AF700, 53-6.7, 100730, BioLegend), γδTCR (APC, GL3, 118115, BioLegend), B220 (PB, RA3-6B2, 103230, BioLegend), and NK1.1 (FITC, PK136, 108706, BioLegend) antibodies were used for the analysis of mature cells in the spleen and lymph nodes. Anti-CD45.1 (FITC, A20, 110706, BioLegend) and CD45.2 (APC, 104, 109814, BioLegend) antibodies were used for the analyses of donor chimerism in the bone marrow transplantation assay. All antibodies used in this study are listed in **Supplementary Table 1**. Dead cells were excluded by 4,6-diamidino-2- phenylindole (DAPI) staining.

For cell sorting, DN thymocytes were purified by negative selection using a magnetic bead/column system (CD4 (L3T4) MicroBeads, 130-117-043; CD8a (Ly-2) MicroBeads, 130- 117-044; LD Columns, 130-042-901; all purchased from Miltenyi Biotec) in accordance to the manufacturer’s instructions. The pre-enriched DN cells were further stained as described above to sort DN2, DN3 or DN4 cells. Dead cells were excluded by DAPI staining.

Flow cytometry data were collected using BD FACS Diva 8 on a BD LSR II or BD Fortessa analyzer. The cell-sorting experiments were performed on a FACS Aria II cell sorter (BD). The acquired data were analyzed using the FlowJo 10 software.

### Apoptosis assay

For Annexin V staining, thymocytes were first stained with surface makers and then stained with Annexin V antibody for 30 min at room temperature in Annexin V binding buffer (TONBO, TNB-5000-L050). The cells were resuspended in Annexin V binding buffer with 1μg/ml DAPI for analysis.

### Cell Proliferation Assay

For cell cycle analysis, thymocytes were stained with surface markers followed by fixation and permeabilization with eBioscience Transcription Factor Staining Buffer Set (Life Technologies, 00-5523-00). The cells were then stained with anti-Ki-67 (FITC, 16A8, 652409, Biolegend) and DAPI for cell cycle analysis. For BrdU staining, total thymocytes were stained with surface markers to define each subset, followed by intracellular staining using the BrdU staining kit according to the manufacturer’s instructions (BD Biosciences, 559619).

### Cell culture

OP9 bone marrow stromal cells expressing the Notch ligand DL-1 (OP9-DL1) were kindly provided by Dr. Juan Carlos Zúñiga-Pflücker (University of Toronto). The OP9-DL1 cells were cultured and maintained in αMEM medium, supplemented with 10% FBS (Gibco), penicillin (100 μg/mL) and streptomycin (100 U/mL) (Invitrogen). For thymocytes co-cultures, the sorted DN2 or DN3 cells were plated onto confluent OP9-DL1 monolayers (70–80% confluent) with addition of 5 ng/ml recombinant murine interleukin-7 (PeproTech) and 5 ng/ml Flt3L (PeproTech). Cells were harvested at different time points, filtered through 40μm cell strainer, and stained with antibodies including ani-CD45 V450 (Tonbo, 75-0451-U100), anti-CD4 BV650 (Biolegend, 100546), anti-CD8a AF700 (Biolegend, 100730), anti-CD44 FITC (Biolegend, 103006), or anti-CD25 APC (Biolegend, 101910), followed by staining with Annexin V PE (Biolegend, 640908) and DAPI (4,6-diamidino-2-phenylindole, Invitrogen, D1306). EL4 cells (ATCC-TIB39) were cultured in DMEM with 10% FBS (Gibco), penicillin (100 μg/mL) and streptomycin (100 U/mL) (Invitrogen).

### Immunohistochemical Staining

Thymus was fixed in fresh 4% paraformaldehyde for 24 hours and stored in 70% ethanol until paraffin embedding. Hematoxylin and eosin staining were performed on 5 μm–thick paraffin sections. For cleaved caspase-3 IHC, the anti-cleaved caspase-3 antibody (1:50, Cell Signaling Technology #9661) was used. Slides were incubated with Envision Labelled Polymer-HRP Anti-Rabbit (Dako, K4002) for 30 minutes. Sections were developed with DAB+ solution (Dako, K3468) and counterstained with Harris Hematoxylin. Imaging analysis was performed with ImageJ (FIJI) to automatically count the labeled cells in each region, the data were shown as number of positive cells per region.

### RNA extraction and Quantitative real-time PCR

Thymocytes were sorted directly into TRIzol LS reagent (Invitrogen, 10296010). Total RNA was extracted according to the manufacturer’s instructions. Total RNA was reverse-transcribed using High-Capacity cDNA Reverse Transcription Kit (Thermo Fisher Scientific, 4368813).

Quantitative real-time PCR was performed using PowerUp™ SYBR™ Green Master Mix (Thermo Fisher Scientific, A25778) on QuantStudio 6 real-time PCR system (Applied Biosystems). The primer sequences are listed in **Supplementary Table 2**.

### Western blot analysis

Western blot was performed as described previously (Xu et al., 2020). Approximately 5 × 10^5^ DN3 cells were sorted directly into 250 μl PBS containing 20% trichoracetic acid (TCA). The concentration of TCA was adjusted to 10% after sorting and cells were incubated on ice for 30 min before centrifugation at 13,000 rpm for 10 min at 4 °C. Precipitates were washed twice with pure acetone (Fisher scientific, A18-4) and solubilized in 9 M urea, 2% Triton X- 100, and 1% dithiothreitol (DTT) in 1 x LDS sample buffer (Invitorgen, NP0007). Samples were separated on NuPAGE 4–12% Bis-Tris protein gels (Invitrogen, NP0336BOX) and transferred to PVDF membrane (Millipore). The blots were incubated with primary antibodies overnight at 4 °C and then with secondary antibodies. Blots were developed with the SuperSignal West Femto chemiluminescence kit (Thermo Scientific, 34096). Antibodies and reagents used are in **Supplementary Table 1 and 3**. The original western blot images are in source data files.

### RNA-seq and analysis

DN3 thymocytes were directly sorted into TRIzol LS (ThermoFisher, cat. 10296028) and RNA was extracted following the standard protocol. The cDNA libraries were prepared using Truseq Stranded mRNA Kit (Illumina, California, USA # 20020594). Sequencing was performed on Illumina HiSeq 2000 (Illumina, California, USA), 150 bp paired end. Quality control was performed using FastQC. Raw reads were aligned to mouse genome GRC38 using STAR (2.5.2b) and counts for each protein-coding gene were obtained with HTSeq (2.7). DESeq2 package (1.26.0) in R (3.6.1) was used to perform analysis of differential gene expression.

Upregulated genes (with adjusted p value < 0.05, log2 fold change > 0.59) were selected for Gene Ontology (GO) analysis using Enrichr (https://maayanlab.cloud/Enrichr/). We further performed Gene Set Enrichment Analysis (GSEA) using fgsea package (1.11.2) with padj- preranked gene lists and mouse gene set collection from Bader Lab. Heatmaps were generated with pheatmap (1.0.12).

### Chromatin Immunoprecipitation (ChIP) assay

EL4 cells were treated with PBS or 5 μg/ml recombinant mouse DLL4 (BioLegend, #776706) for 24 hours before crosslinked with 1% formaldehyde for 10 minutes at room temperature.

Reaction was quenched with 125 mM glycine. ChIP was performed as previously described (Zhao et al., 2018) with NOTCH1 antibody (Abcam, ab27526), RBPJ antibody (Cell Signaling Technology, #5313), or normal rabbit IgG (Cell Signaling Technology, #2729). The sequences of all ChIP primers are listed in **Supplemental Table 2**.

### Luciferase assay

The firefly luciferase reporter for *Sel1l* or *Hrd1* (*Syvn1*) promoter was constructed by cloning the genomic region into the *Mlu*I and *Xho*I sites or the *Mlu*I and *Hind*III sites in the pGL-3 basic vector (Promega), respectively. Mutations were made by overlap extension polymerase chain reaction as previously described (Bryksin and Matsumura, 2010). All constructs were verified by DNA sequencing. The sequences of all primers are listed in **Supplemental Table 2**. 293T cells were transfected with *Sel1l* or *Hrd1* promoter constructs, pRL-PGK (Promega) and 3xFlag- NICD1 (Addgene, #20183) or empty using Lipofectamine 3000 (Invitrogen, L3000015). Cell lysates were collected 48 hours after transfection, and luciferase activities were analyzed using the dual-luciferase reporter assay system (E1910, Promega). pRL-PGK, which expresses Renilla luciferase, was used as the internal control for adjustment of discrepancies in transfection and harvest efficiencies. EL4 cells were transfected with *Sel1l* or *Hrd1* promoter constructs and pRL- PGK (Promega) using Lipofectamine 3000 (Invitrogen, L3000015). Cells were incubated with PBS or 5 μg/ml recombinant mouse DLL4 (BioLegend, #776706) for 24 hours before analysis.

### V(D)J recombination assay

5 × 10^5^ DN3 thymocytes were sorted from control (Ctrl, *Sel1l^flox/flox^*) or *Sel1l^CD2^* (*Sel1l^flox/flox^* ; *hCD2-iCre*) thymus and subjected to genomic DNA isolation using PureLink® Genomic DNA Mini Kit (K1820-02) according to the manufacturer’s instructions. The amplification of eF-1 fragment was used as input control. The primers used for Vβ5-Jβ2, Vβ8- Jβ2, Vβ11-Jβ2, and eF1 are listed in **Supplementary Table 2**. PCR products were resolved by 2% agarose gel electrophoresis.

### ER tracker staining

Sorted DN3 thymocytes were washed with PBS, incubated with 1μM ER-Tracker Green (Thermo Fisher, E34251) in PBS for 15 min at 37 °C. The cells were then washed and resuspended in PBS, and analyzed by flow cytometry.

### Tetraphenylethene maleimide (TMI) staining

5x10^6^ thymocytes were stained for cell surface markers as described above. After surface markers staining, cells were washed twice with PBS. Tetraphenylethene maleimide (TMI; 2 mM in DMSO) was diluted in PBS to reach 50 μM final concentration and stained samples for 30 minutes at 37 °C. Samples were washed once with PBS and analyzed by flow cytometry.

### *In vivo* measurement of protein synthesis

O-propargyl-puromycin (OPP; MedChem Source LLP) stock dissolved in 10% DMSO/PBS was diluted in PBS for intraperitoneal injection at 50mg/kg. Mice were weighed, individually injected with OPP or vehicle, and euthanized one hour after injection. Thymuses were immediately harvested. Thymocytes were isolated and stained for surface antigens and viability dyes, which was followed by fixation and permeabilization according to the Click-iT Plus OPP Alexa Fluor 647 Protein Synthesis Assay Kit (ThermoFisher Scientific). Briefly, after incubation cells were subsequently fixed in 4% paraformaldehyde, permeabilized with 0.5% Triton-X100, then incubated with the AF647 reaction cocktail. Samples were acquired using a BD LSRFortessa and analyzed using FlowJo (Becton Dickinson) as per flow cytometry methods.

### Protein aggregation detection assay

The PROTEOSTAT Aggresome Detection kit (Enzo Life Sciences, ENZ-51035-0025) was used to detect protein aggregates in freshly sorted DN3 thymocytes according to the manufacturer’s instructions. DN3 thymocytes were fixed, permeabilized and incubated with PROTEOSTAT dye (1:10,000 dilution) for 30 min at room temperature. Nuclei were counterstained with DAPI. Samples stained with DAPI only were used as negative controls. Images (16-bit greyscale TIFFs) were analyzed using CellProfiler v2.2. In brief, the DAPI channel images were first smoother with a median filter and nuclei identified with automatic thresholding and a fixed diameter.

Nuclei touching the border of the image are eliminated. Touching nuclei are separated with a watershed algorithm. Then, cell boundaries were identified by watershed gradient based on the dye signal, using nuclei as a seed. Metrics were extracted from the cell, cytoplasm and nuclear compartments.

### TUNEL staining

TUNEL staining was performed on paraffin-embedded tissue sections using the In Situ Cell Death Detection Kit (catalog 11684795910, Roche) following the manufacturer’s instructions.

Sections were counterstained with DAPI, and images were captured under fluorescence microscope. Tissue sections incubated with TUNEL reaction buffer without dTd enzyme served as negative controls. Tissue sections treated with DNase I served as positive controls. The quantification of TUNEL^+^ cells was performed with ImageJ (FIJI) to automatically count the labeled cells in each region. The data were presented as number of positive cells per region.

### Statistics and reproducibility

Data are expressed as the mean ± s.d. or mean ± s.e.m. as indicated in the figure legends; *n* is the number of independent biological replicates, unless specifically indicated otherwise in the figure legend. The respective *n* values are shown in the figure legends. The mice used for bone marrow transplantation were randomized and no blinding protocol was used. No statistical method was used to pre-determine the sample sizes. The results were quantified using GraphPad Prism 8. *P* values were generated using two-tailed unpaired/paired Student’s t-tests as indicated.

### Study approval

All protocols described in this study were approved by the Baylor College of Medicine Institutional Animal Care and Use Committee or Case Western Reserve University Institutional Animal Care and Use Committee.

### Data availability

Sequencing data have been deposited in GEO under accession code GSE173993. All the numerical data and the original western blots supporting the findings of this study are provided in the source data submitted with the manuscript.

---

**Figure 1 - figure supplement 1.**
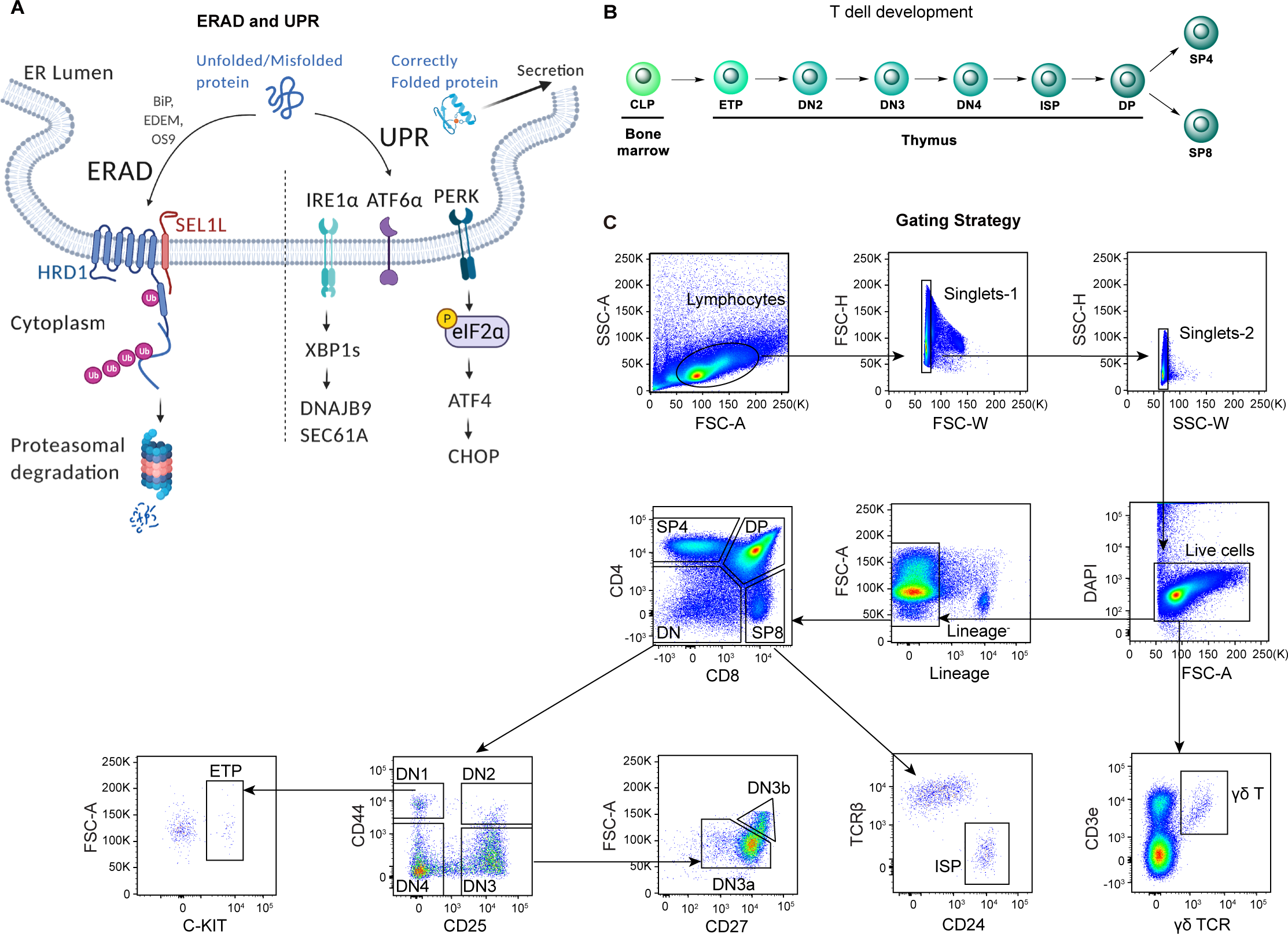
Diagrams and representative flow cytometry gates used in this study. **(A)**, Diagram showing two ER quality control machineries: ERAD and UPR. The E3 ubiquitin ligase HRD1 and its adaptor protein SEL1L is the most conserved ERAD complex in mammals. While correctly folded proteins exit the ER, misfolded proteins in the ER are recruited to the SEL1L-HRD1 complex through ER chaperones (such as BiP, EDEM, and OS9), and then retrotranslocated into the cytosol, ubiquitinated and degraded by the proteasome. Failure to clear the misfolded or unfolded proteins in the ER activates the UPR signaling through three ER stress sensors IRE1*α*, ATF6 and PERK. Upon activation, IRE1*α* oligomerizes and undergoes *trans*- autophosphorylation to activate its RNase domain, resulting in the removal of 26 nucleotides from unspliced *XBP1* (*XBP1u*) mRNA to produce mature, spliced *XBP1* (*XBP1s*) mRNA. PERK is a serine-threonine kinase. ER stress induces PERK-dependent eIF2*a* phosphorylation and subsequent increased cap-independent translation of ATF4 and induction of CHOP. **(B)**, Schematic diagram of T-cell development in the thymus. CLP: common lymphoid progenitors; ETP: early T lineage precursor (Lin^-^ CD4^-^ CD8^-^ CD44^+^ CD25^-^ CD117^+^); DN2: double negative 2 thymocytes (Lin^-^ CD4^-^ CD8^-^ CD44^+^ CD25^+^); DN3: double negative 3 thymocytes (Lin^-^ CD4^-^ CD8^-^ CD44^-^ CD25^+^); DN4: double negative 4 thymocytes (Lin^-^ CD4^-^ CD8^-^ CD44^-^ CD25^-^); ISP: immature single-positive thymocytes (Lin^-^CD8^+^CD24^+^TCRβ^-^); DP: double positive thymocytes (Lin^-^ CD4^+^ CD8^+^); SP4: CD4 single positive thymocytes (Lin^-^ CD4^+^ CD8^-^); SP8: CD8 single positive thymocytes (Lin^-^ CD4^-^ CD8^+^). **(C)**, Representative pseudocolor plots showing the gating strategy to identify different thymocyte subsets in the thymus.

**Figure 2 - figure supplement 1.**
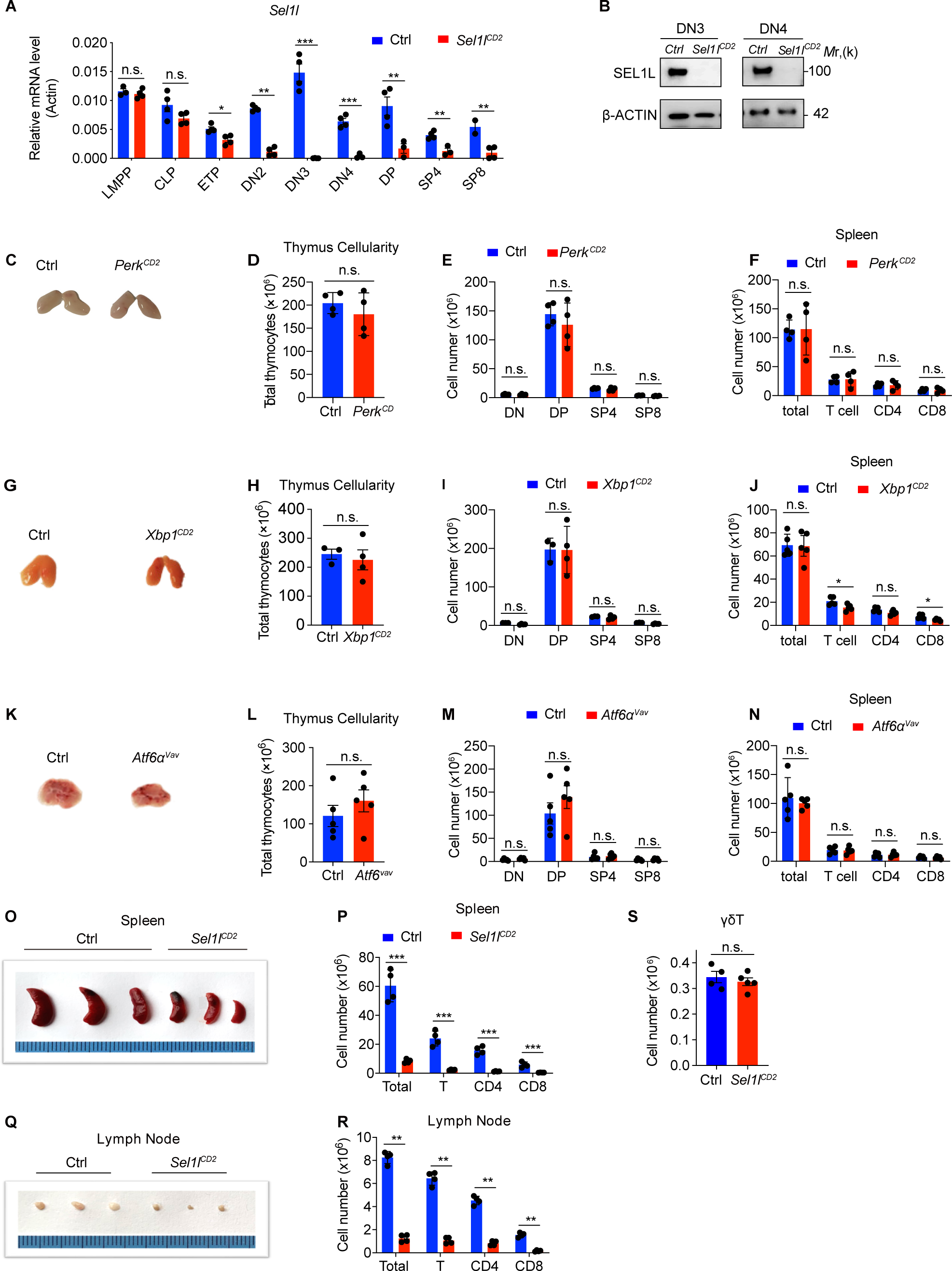
UPR is dispensible for αβ T cell development. **(A)**, Quantitative RT-PCR analysis of *Sel1l* expression in murine bone marrow progenitors and different thymocyte subsets from control (Ctrl, *Sel1l^flox/flox^*) or *Sel1l^CD2^*-KO (*Sel1l^flox/flox^*; *hCD2- iCre*) mice. Data are presented relative to *Actin*. *n* = 4. LMPP: lymphoid-primed multipotent progenitor. **(B)**, Western blot analysis of SEL1L protein in sorted DN3 and DN4 thymocytes from Ctrl or *Sel1l^CD2^* mice. β-ACTIN was used as loading control. The original western blot images are provided in Figure 2 **- figure supplement 1- source data 1**. **(C-F),** Representative images of thymus (**C**), thymus cellularity (**D**), cell numbers of indicated populations in the thymus (**E**) and peripheral splenocyte numbers of indicated populations (**F**) from age and gender-matched control (Ctrl, *Perk^flox/flox^*) or *Perk^CD2^-*KO (*Perk^flox/flox^*; *hCD2-iCre*) mice. *n* = 4. **(G-J),** Representative images of thymus (**G**), thymus cellularity (**H**), cell numbers of indicated populations in the thymus (**I**) and peripheral splenocyte numbers of indicated populations (**J**) from age and gender-matched control (Ctrl, *Xbp1^flox/flox^*) or *Xbp1^CD2^-*KO (*Xbp1^flox/flox^*; *hCD2- iCre*) mice. *n* = 3-4. **(K-N),** Representative images of thymus (**K**), thymus cellularity (**L**), cell numbers of indicated populations in the thymus (**M**) and peripheral splenocyte numbers of indicated populations (**N**) from age and gender-matched control (Ctrl, *Atf6^flox/flox^*) and *Atf6^Vav^-*KO (*Atf6^flox/flox^*; *Vav-iCre*) mice. *n* = 5. **(O)**, Images of spleen from 6-8 week-old control (Ctrl, *Sel1l^flox/flox^*) and *Sel1l^CD2^-*KO (*Sel1l^flox/flox^* ; *hCD2-iCre*) mice. **(P)**, Quantification of cell numbers of the indicated populations in the spleen of 6-8 week-old control (Ctrl, *Sel1l^flox/flox^*) and *Sel1l^CD2^-*KO (*Sel1l^flox/flox^* ; *hCD2-iCre*) mice. **(Q)**, Images of the inguinal (left) lymph nodes from 6-8 week-old control (Ctrl, *Sel1l^flox/flox^*) and *Sel1l^CD2^-*KO (*Sel1l^flox/flox^* ; *hCD2-iCre*) mice. **(R)**, Quantification of cell numbers in the lymph nodes of 6-8 week-old control (Ctrl, *Sel1l^flox/flox^*) and *Sel1l^CD2^-*KO (*Sel1l^flox/flox^* ; *hCD2-iCre*) mice. *n* = 4. **(S)**, Quantification of cell numbers of *γδ* T cells from 6-8 week-old control (Ctrl, *Sel1l^flox/flox^*) and *Sel1l^CD2^-*KO (*Sel1l^flox/flox^* ; *hCD2-iCre*) mice. Ctrl: *n*=4. *Sel1l^CD2^-*KO: *n*=5. Data are representative of three independent experiments and are shown as mean ± s.d. Two- tailed Student’s t-tests (**A, D-F, H-J, L-N, P, R, S**) was used to calculate *P* values. n.s., not significant, \**P* < 0.05, \*\**P* < 0.01, \*\*\**P* < 0.001, \*\*\*\**P* < 0.0001.

**Figure 2 - figure supplement 2.**
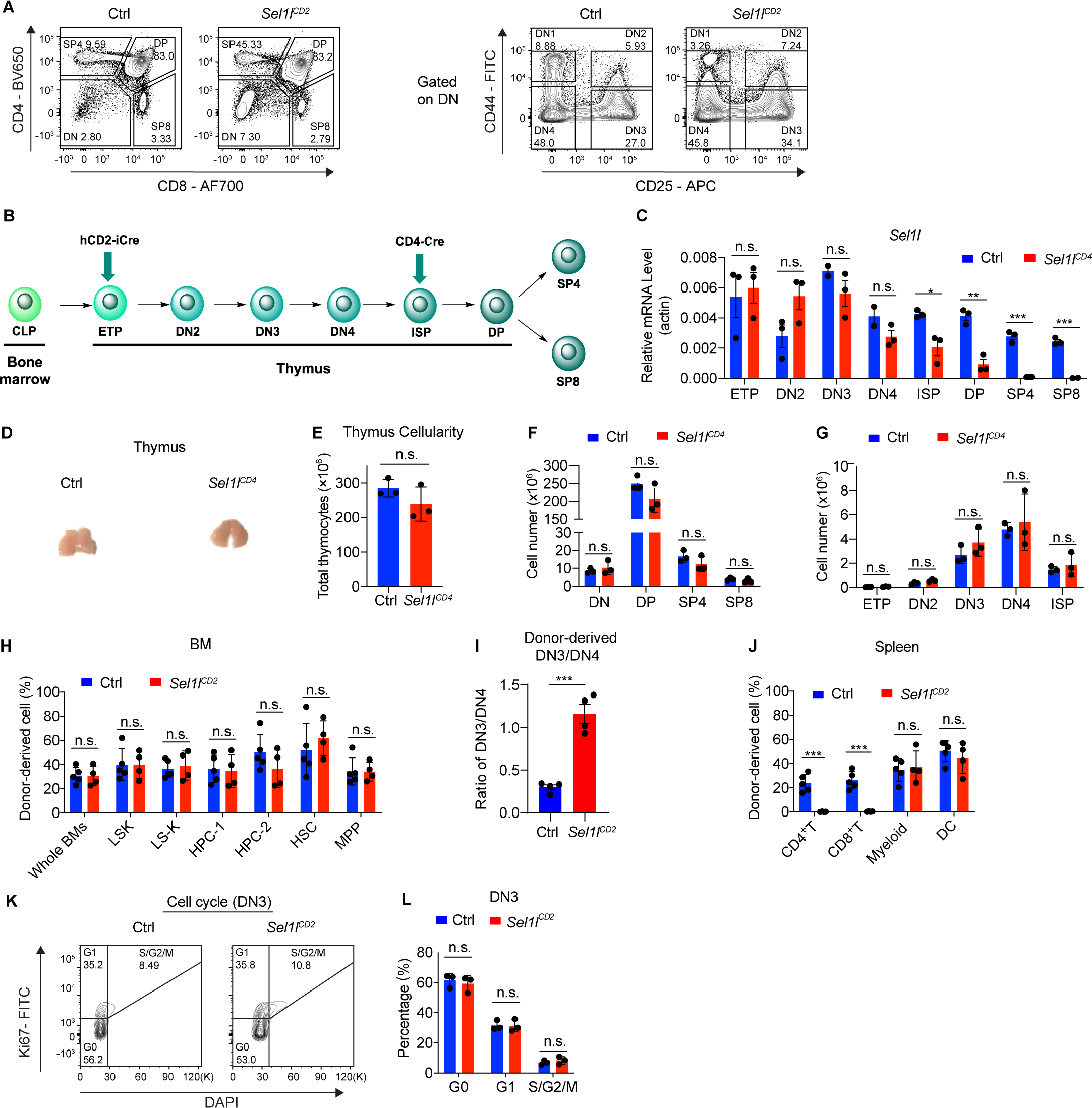
SEL1L is required for DN to DP thymocyte transition following β selection. **(A)**, Representative flow cytometry plots of different thymocyte subsets in control (Ctrl, *Sel1l^flox/flox^*) and *Sel1l^CD2^-*KO (*Sel1l^flox/flox^* ; *hCD2-iCre*) mice. **(B)**, Diagram showing different stages of *hCD2-iCre* and *CD4-iCre* initiated gene depletion during T cell development. **(C)**, Quantitative RT-PCR analysis of *Sel1l* in different thymocyte subsets from control (Ctrl, *Sel1l^flox/flox^*) and *Sel1l^CD4^*-KO (*Sel1l^flox/flox^* ; *CD4-iCre*) mice. Data are presented relative to *Actin*. *n* = 3. **(D and E)**, Representative images of thymus (**D**) and quantification of thymus cellularity (**E**) in 6-8 week-old control (Ctrl, *Sel1l^flox/flox^*) and *Sel1l^CD4^*-KO (*Sel1l^flox/flox^* ; *CD4-iCre*) mice. *n* = 3. **(F and G)**, Quantification of cell numbers of different thymocyte subsets from control (Ctrl, *Sel1l^flox/flox^*) and *Sel1l^CD4^*-KO (*Sel1l^flox/flox^* ; *CD4-iCre*) mice. *n*=3. **(H)**. Percentage of Ctrl or *Sel1l^CD2^-*KO donor-derived progenitors in the bone marrow of recipient mice 14 weeks after transplantation. *n* =4-5. **(I)**, Quantification of Ctrl or *Sel1l^CD2^-*KO donor-derived DN3/DN4 ratio. *n* =4-5. **(J)**, Percentage of Ctrl or *Sel1l^CD2^-*KO donor-derived CD4^+^ T cells, CD8^+^ T cells, myeloid cells, and dendritic cells (DC) in the spleen of recipient mice 14 weeks after transplantation. *n* = 4-5. **(K and L),** Cell cycle analysis of DN3 thymocytes in 6-week-old control (Ctrl) and *Sel1l^CD2^* mice using Ki67 and DAPI. Representative flow cytometry plots (**K**) and quantification (**L**) are shown. *n* = 3. Data are shown as mean ± s.d. The statistical significance was calculated by two-tailed unpaired t-test (**C, E-J**) or Two-way ANOVA with Bonferroni test (**L**). n.s., not significant, \**P* < 0.05, \*\**P* < 0.01, \*\*\**P* < 0.001, \*\*\*\**P* < 0.0001.

**Figure 4 - figure supplement 1.**
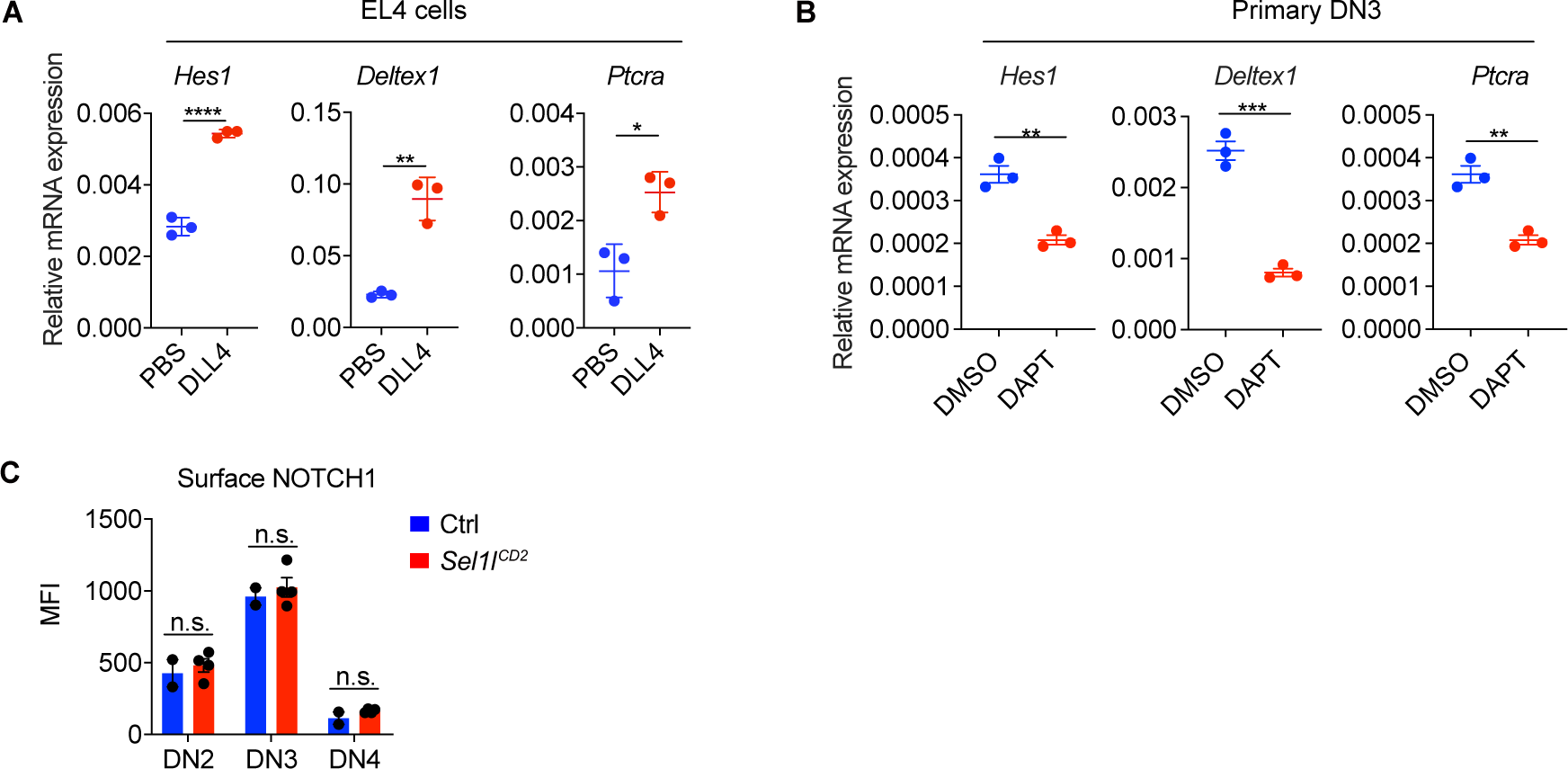
Notch signal regulates ERAD genes expression. **(A)**, Quantitative RT-PCR analysis of Notch target genes expression in EL4 cells after stimulation with 5 μg/ml Delta ligand 4 (DLL4) for 24h. Data are presented relative to *Actin*. *n*=3. **(B)**, Quantitative RT-PCR analysis of Notch target genes expression in primary DN3 thymocytes treated with 2mM *γ*-secretase inhibitor DAPT for 5 hours. Data are presented relative to *Actin*. *n*=3. **(C)**, Expression of Notch1 on cell surface of different thymocyte subsets from control (Ctrl, *Sel1l^flox/flox^*) and *Sel1l^CD2^-*KO (*Sel1l^flox/flox^* ; *hCD2-iCre*) mice. Ctrl: *n* = 2. *Sel1l^CD2^*-KO: *n*=4. Data are shown as mean ± s.d. Two-tailed Student’s t-tests (**A**-**C**) was used to calculate *P* values. n.s., not significant, \**P* < 0.05, \*\**P* < 0.01, \*\*\**P* < 0.001, \*\*\*\**P* < 0.0001.

**Figure 5 - figure supplement 1.**
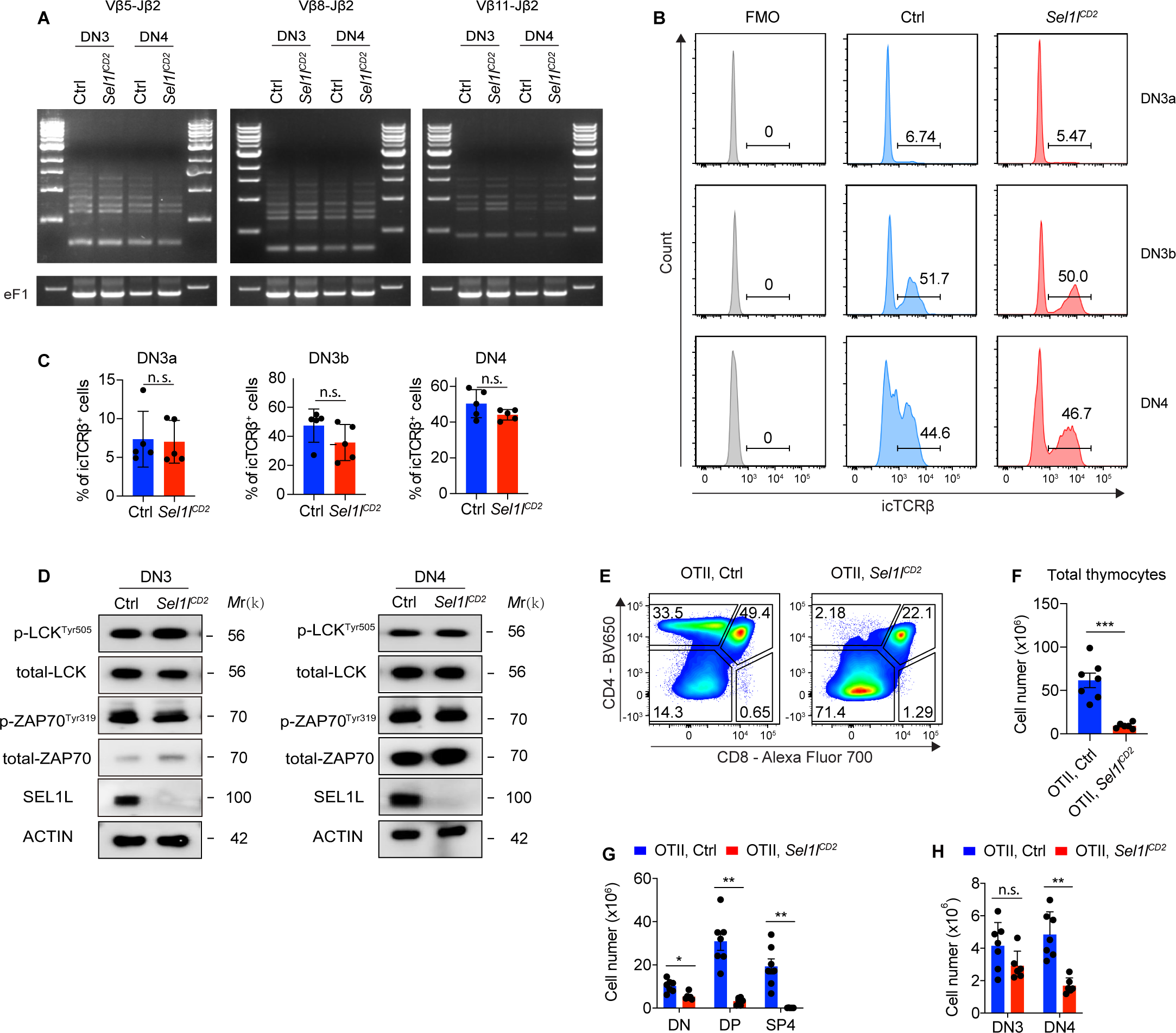
SEL1L is not required for TCRβ gene rearrangement and pre-TCR signaling. **(A)**, PCR analysis of *Vβ5-Jβ2*, *Vβ8-Jβ2* and *Vβ11-Jβ2* gene rearrangements using genomic DNA of DN3 and DN4 thymocytes sorted from control (Ctrl, *Sel1l^flox/flox^*) or *Sel1l^CD2^-*KO (*Sel1l^flox/flox^* ; *hCD2-iCre*) mice. The original gel images are provided in Figure 5 **- figure supplement 1- source data 1.** **(B and C),** Representative flow cytometry plots (**B)** and quantification (**C**) of intracellular TCRβ positive cells in DN3a, DN3b and DN4 thymocytes from Ctrl or *Sel1l^CD2^-*KO mice. *n* = 5. **(D)**, Western blot analysis of the expression of proteins involved in pre-TCR signaling in primary DN3 and DN4 thymocytes sorted from Ctrl or *Sel1l^CD2^* mice. β-ACTIN was used as loading control. The original western blot images are provided in Figure 5 **- figure supplement 1- source data 2.** **(E-H),** Representative pseudocolor plots (**E**), quantification of total thymocytes (**F**) and cell numbers of indicated populations (**G and H**) from OT-II.Ctrl (*OT-II*; *Sel1l^flox/flox^*) or OT- II.*Sel1l^CD2^* (*OT-II*; *Sel1l^flox/flox^* ; *hCD2-iCre*) mice. OT-II.Ctrl: *n* = 7. OT-II.*Sel1l^CD2^*: *n* = 6. Data are shown as mean ± s.d. Two-tailed Student’s t-tests (**C, F-H**) was used to calculate *P* values. n.s., not significant, \**P* < 0.05, \*\**P* < 0.01, \*\*\**P* < 0.001, \*\*\*\**P* < 0.0001.

**Figure 5 - figure supplement 2.**
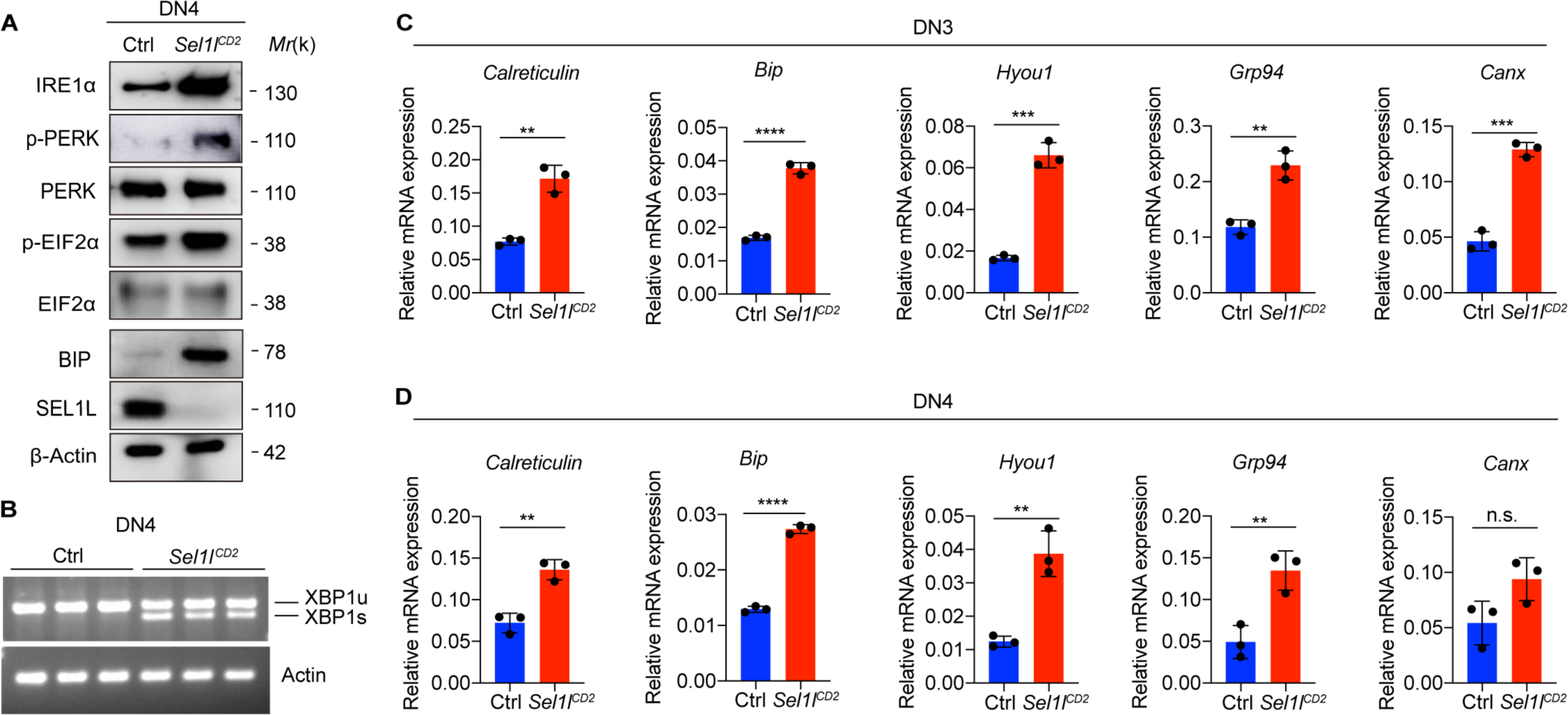
*Sel1l* knockout induces ER stress. **(A),** Western blot analysis of UPR pathway markers in primary DN4 thymocytes sorted from 6- week-old Ctrl or *Sel1l^CD2^-*KO mice. β-ACTIN was used as loading control. The original western blot images are provided in Figure 5 **- figure supplement 2- source data 1.** **(B),** PCR analysis of XBP1-splicing in DN4 thymocytes sorted from Ctrl or *Sel1l^CD2^* mice. Xbp1u: Unspliced Xbp1; Xbp1s: Spliced Xbp1. β-ACTIN was used as loading control. The original gel images are provided in Figure 5 **- figure supplement 2- source data 2.** **(C and D)**, Quantitative RT-PCR analysis of ER chaperone genes expression in DN3 (**C**) and DN4 (**D**) thymocytes sorted from 6-week-old Ctrl or *Sel1l^CD2^*-KO mice. Data are presented relative to *Actin*. *n* = 3. Data are shown as mean ± s.d. Two-tailed Student’s t-tests was used to calculate *P* values. n.s., not significant, \**P* < 0.05, \*\**P* < 0.01, \*\*\**P* < 0.001, \*\*\*\**P* < 0.0001.

**Figure 6 - figure supplement 1.**
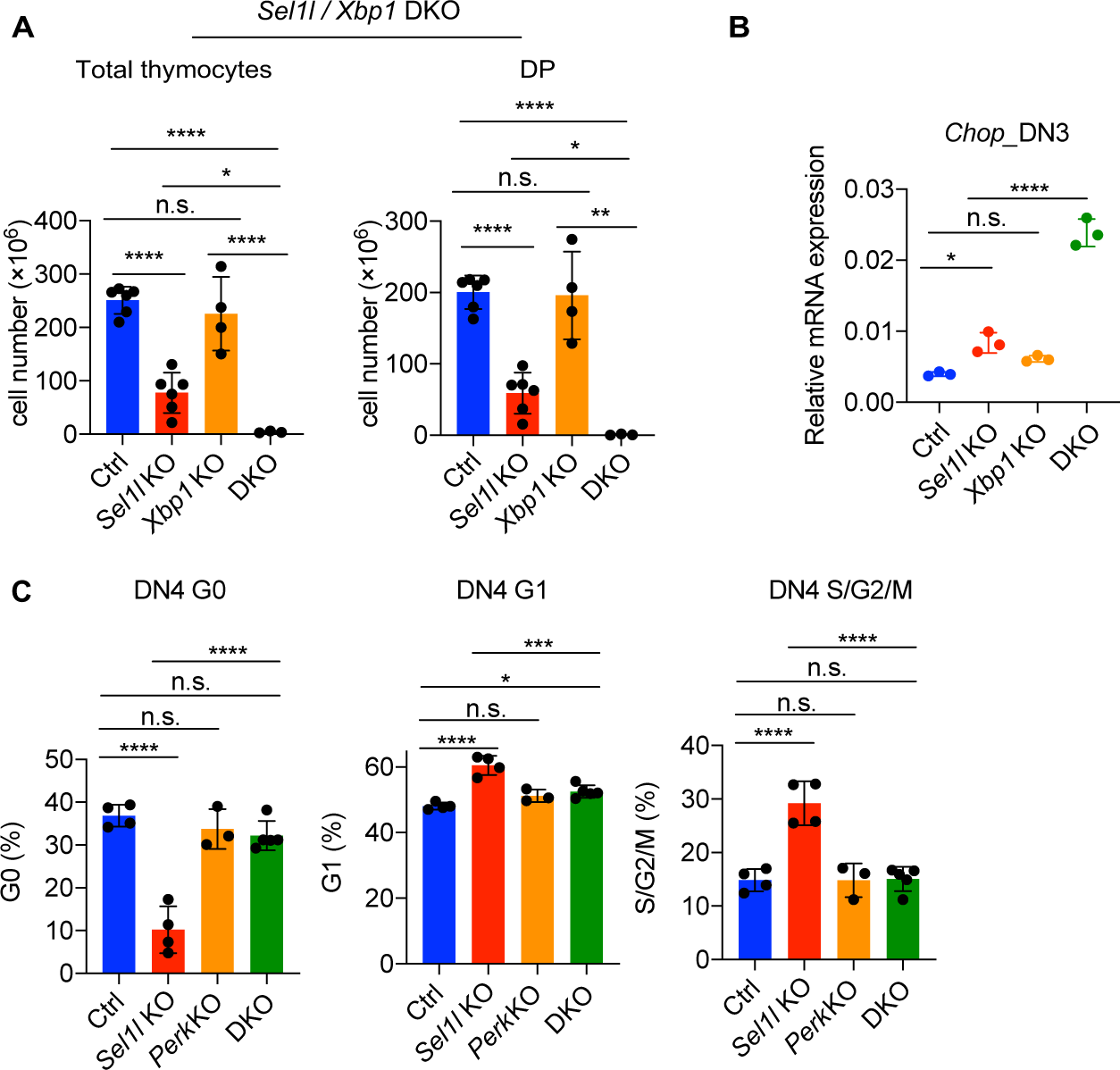
XBP1 functions as a compensatory adaptative mechanism in *Sel1l*-KO mouse. **(A)**, Quantification of total cellularity and DP cell numbers of 6-8 week-old gender-matched control (Ctrl, *Sel1l^flox/flox^*), *Sel1l-*KO (*Sel1l^flox/flox^* ; *hCD2-iCre*), *Xbp1-*KO (*Xbp1 ^flox/flox^* ; *hCD2- iCre*), and *Sel1l*/*Xbp1* double knockout (DKO, *Sel1l^flox/flox^; Xbp1^flox/flox^* ; *hCD2-iCre*) mice. *n* = 3- 6/each group. **(B),** Quantitative RT-PCR analysis *Chop* expression in DN3 thymocytes sorted from mice with indicated genotype. Data are presented relative to *Actin*. **(C)**, Cell cycle analysis of DN4 thymocytes from age (6-week-old) and gender-matched control (Ctrl, *Sel1l^flox/flox^*), *Sel1l*-KO (*Sel1l^flox/flox^* ; *hCD2-iCre*), *Perk-*KO (*Perk^flox/flox^* ; *hCD2-iCre*)and *Sel1l*/*Perk* double knockout (DKO. *Sel1l^flox/flox^; Perk^flox/flox^* ; *hCD2-iCre*) mice. *n* = 3-5 each group. Data are shown as mean ± s.d. The statistical significance was calculated by two-tailed unpaired t-test (**A, B**) or One-way ANOVA with Turkey test (**C**). n.s., not significant, \**P* < 0.05, \*\**P* < 0.01, \*\*\**P* < 0.001, \*\*\*\**P* < 0.0001.

## Notes

### Competing Interest Statement

The authors have declared no competing interest.

